# Temporally Dynamic Waves of Selection on Archaic Variation in Peruvians

**DOI:** 10.64898/2026.11.13.766243

**Authors:** Susanna Gutierrez, Autumn Allemon, Elizabeth Meyer, Xinjun Zhang

## Abstract

Introgression introduced deeply diverged archaic variation into modern human populations and contributed to local adaptation. In Indigenous American populations, however, the landscape of adaptive introgression remains poorly resolved. Genome-wide scans have reported numerous candidate loci, but complex demographic histories and recent admixture make genuine signals difficult to distinguish from false positives. Here, we systematically re-evaluated previously reported candidates of adaptive introgression in Peruvians using a multilayered framework and refined more than 150 putative signals to 25 high-confidence candidate loci with concordant evidence of Neanderthal- or Denisovan-like ancestry and positive selection. Using ancestral recombination graph (ARG) approaches, we inferred selection onset windows for 21 loci. These onsets were non-uniformly distributed across Peruvian population history, revealing temporally dynamic selection without robust evidence for a fixed number of discrete waves. Together with pre-Columbian ancient DNA evidence, the inferred histories support the inheritance of the candidate adaptive archaic haplotypes through ancestral Native American populations rather than first introduction through post-Columbian admixture. Functional analyses identified network-level signals consistent with polygenic selection in lipid metabolic and immune-related pathways, elevated regulatory effects predicted across archaic haplotypes, and occasional coding variation of potential functional relevance, including a missense variant at the immune-related gene *CXCR4*. Our results show that archaic introgressed variation remained evolutionarily important and available to selection long after introgression, providing a reservoir of genetic variation that could be repeatedly recruited by selection at different points in American population history.

**Significance Statement:** Archaic variants reached the Americas through populations already carrying Neanderthal and Denisovan ancestry, offering an opportunity to study adaptation far from the original contact zone. Scans of American populations have reported more putative adaptive introgression loci than realistically expected, partly because the complex demographic histories make true signals difficult to distinguish from false positives. Reconstructing selection histories at credible loci may in turn help constrain that history. Here we identify 25 high-confidence candidate regions in Peruvians, among which 21 have inferred selection onsets predating European contact, and archaic-associated alleles from all 25 are present in pre-Columbian American ancient genomes. Functional analyses find pathway-level selection signals, predicted regulatory effects, and occasional evolutionarily interesting missense variants, including at the immune-related gene *CXCR4*.

## Introduction

Archaic introgression introduced deeply divergent genetic variation into the modern human gene pool (1-3). Much of this variation was selectively neutral or deleterious and subsequently declined in frequency over time through negative selection and genetic drift, while recombination broke introgressed segments into smaller haplotypes (4-6). In some cases, however, introgressed haplotypes conferred a fitness advantage and rose to higher frequencies in modern humans, a process known as adaptive introgression (7, 8). Classic examples of adaptive introgression include the Denisovan-like haplotype at the *EPAS1* gene associated with high-altitude adaptation in Tibetans (9) and archaic variation at the *BNC2* gene associated with pigmentation phenotypes in Europeans (10-12). These cases establish archaic introgression as an important source of functional variation in modern humans.

The contribution of archaic introgression to adaptation in the Americas is less clear. Archaic ancestry reached the Americas indirectly through populations that had admixed with Neanderthals and Denisovans before the peopling of the continent. Genome-wide scans reported many candidate loci of adaptive introgression in American populations (13-16), while only a few, including *TBX15*/*WARS2* (17) and *MUC19* (18) have been examined in sufficient detail to resolve their evolutionary histories and potential phenotypes. Therefore, the problem with studying adaptive introgression in the Americas is not a lack of candidate loci, but an excess of putative candidates relative to the number likely to represent genuine positive selection events.

One reason for this apparent excess is the unusually complex demographic history of American populations. Severe bottleneck events during the peopling of America, subsequent population structure and changes in population size, and recent post-Columbian admixture can all generate patterns that resemble positive selection or obscure genuine signals (19). European admixture can also introduce archaic variants that are common in European populations, complicating the distinction between variation inherited through Native American lineages and more recent European contributions (20). Additionally, because there is no single well-supported demographic model applicable across American populations, methods developed or calibrated primarily under Eurasian demographic histories may produce spurious signals when applied to American populations. These factors could all contribute to elevated false positive rates in adaptive introgression scans.

However, some of the reported candidates are likely genuine, and distinguishing them is particularly valuable. Previous studies have shown that selection on introgressed archaic variation can occur either soon after admixture (21) or later on introgressed variants retained as standing variation (22). Much less is known about how long after introgression variation remains available to selection, or how far populations can move from the original contact zone before archaic introgressed variants become beneficial. The Americas provide an opportunity to address these questions, because the adaptive events on archaic variants in Peruvians necessarily occurred far in time and geography from the original archaic contacts. Reconstructing the selection history of high-confidence archaic adaptive haplotypes can therefore reveal their long-term evolutionary fate and help place selection in the context of the initial peopling of the Americas (23-25), later settlement of South America (26-28), and subsequent ecological and subsistence transitions (29-32). These selection histories may also provide temporal constraints on aspects of American population history that remain difficult to infer from demographic inference alone.

Here in this work, we systematically re-evaluate previously reported adaptive introgression candidates in modern Peruvians, a population with relatively high Indigenous American ancestry among contemporary American populations (33, 34) and numerous previously reported candidate loci (13-15). Starting from over 150 putative regions, we combined archaic ancestry statistics, haplotype structure and positive selection scans to obtain a conservative set of 25 high-confidence adaptive introgression candidate regions. We then use ancestral recombination graphs (ARGs) to reconstruct selection histories for these loci and examined pre-Columbian ancient genomes (20, 35-39) for evidence that archaic alleles were present before European contact. Finally, we ask whether selection onset is temporally structured and associated with major population history transitions, and examine pathway-level, predicted regulatory, and coding consequences of candidate regions.

Our results show that for the 21 regions with resolved selection histories, selection predates European contact and are consistent with inheritance through ancestral Native American population. Selection onset times were non-uniformly distributed across Peruvian population history, and functional analyses further found network-level selection evidence, predicted regulatory consequences, and coding variation in candidate loci including *CXCR4*.

## Results

### Refining the landscape of adaptive introgression in Peruvians

Previous genomic scans reported over 150 putative adaptive introgression signals in Peruvians (PEL in 1000 Genomes Project) (8, 13-15, 40). Rather than maximizing the candidate list, we used a multi-stage framework (**Fig. 1A**) to prioritize loci with concordant evidence for archaic ancestry and positive selection. The well-established Denisovan-derived *EPAS1* haplotype in Tibetans provided a positive control example for this relationship (9, 22). We first screened non-overlapping genomic windows using three empirical outlier criteria: a MaLAdapt score >0.80 (14), a Q95 score (15) above the 95^th^ percentile of the genome-wide distribution, and an average divergence ratio below the 5^th^ percentile of its genome-wide distribution. A high Q95 score indicates a higher frequency of derived alleles shared between modern Peruvian and archaic hominin and absent from an African outgroup, whereas a low divergence ratio identifies unusually similar sequences between Peruvians and archaic genomes. This screening retained 90 candidate windows.

**Fig. 1.**
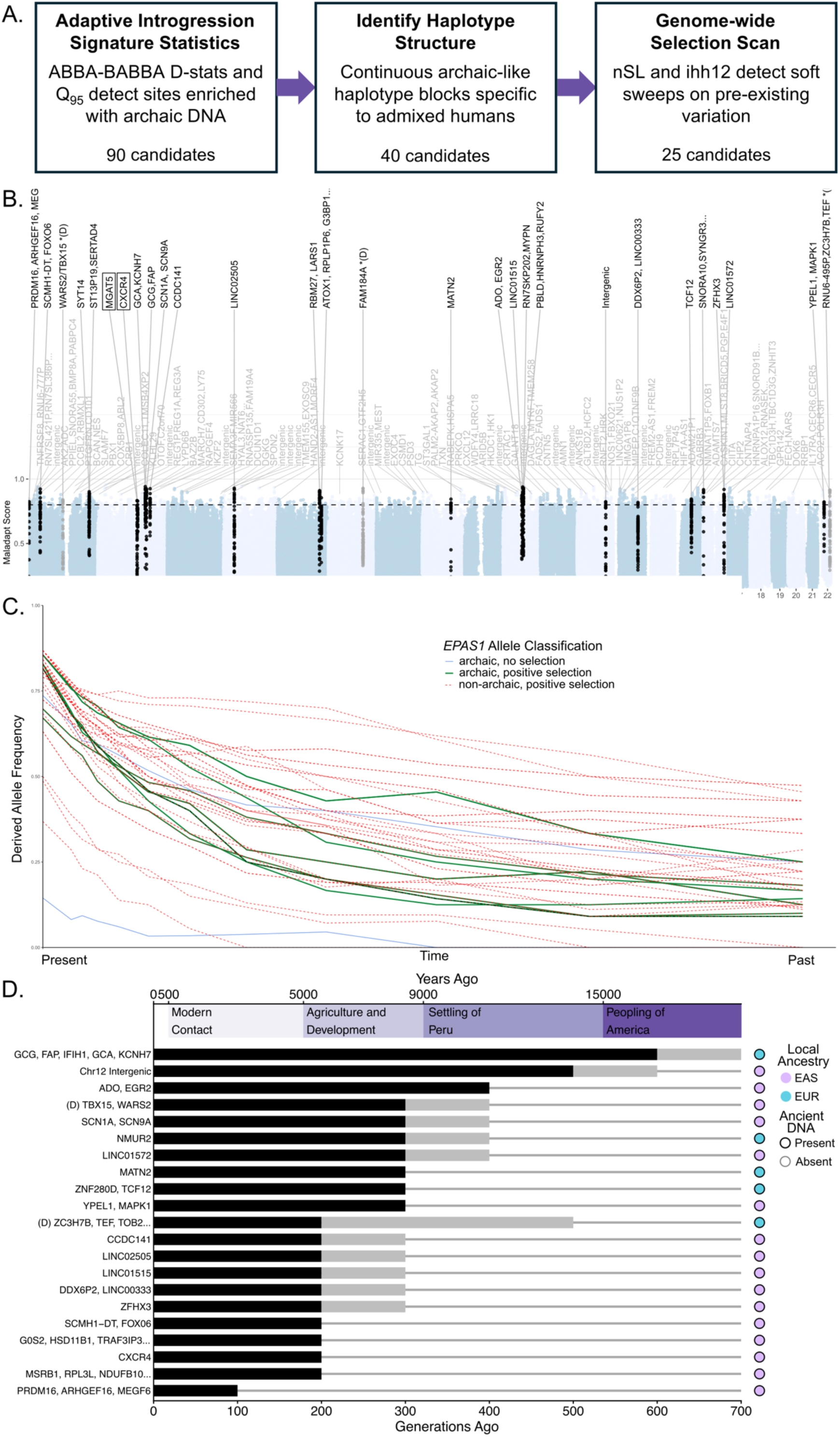
Refined adaptive introgression landscape and reconstructed selection onset times in Peruvians. **A**) Filtering steps used to prioritize archaic-like haplotypes with evidence of selection **B)** Genome-wide distribution of MaLAdapt scores across 50kb windows highlighting candidate genes for adaptive introgression. Gray labels indicate regions previously reported by Zhang et al. 2023. Black labels indicate regions retained here. (D*) indicates Denisovan-like candidates. Boxed gene names indicate candidates not reported by additional studies. Alternating shading indicates chromosomes. **C)** Inferred allele frequency at the EPAS1 positive control region (Chr2: 46500806-46663836; hg19) in Tibetans. Solid lines represent archaic alleles, and dashed lines represent non-archaic alleles with a significant p-value for positive selection. **D)** Selection onset windows in Peruvian candidates. Black bars show intervals with inferred positive selection, and gray bars show uncertainty associated with disagreement between onset criteria. The timeline shows appropriate historical periods with side markers indicate present-day local ancestry assignment and pre-Columbia evidence for archaic alleles.

We next examined the haplotype structure of these regions. Of the 90 candidate windows, 40 contained a distinct cluster of Peruvian haplotypes that was highly similar to Neanderthals or Denisovan, rare or absent in populations with little archaic admixture (such as Yorubans, the YRI population in the 1000 Genomes Studies), and relatively frequent in Peruvians relative to other archaic-admixed modern human populations (e.g. East Asians and Europeans). This identifies archaic haplotypes that rose in frequency after introgression and specifically in the Peruvian lineage. We further tested these regions for positive selection using haplotype-based statistics calculated by Selscan 2.0 (41, 42), including nSL (43), H12 (44), and XP-nSL (45). These statistics are chosen because they are particularly sensitive to soft sweep signatures, which is the pattern we expect from archaic adaptive introgression in Peruvians that are geographically far from the original contact zone between modern humans and archaic hominins. Combining evidence from archaic ancestry statistics, haplotype structure, and positive selection scans yielded 25 high-confidence focal haplotypes, including 23 Neanderthal-like and two Denisovan-like candidates (**Fig. 1B, Table S1, Fig. S1-S2**).

This is an intentionally conservative set. The 25 candidates do not represent every possible case of adaptive introgression in Peruvians, but rather regions with sufficiently concordant evidence to justify closer investigation of their selection histories and potential functional consequences.

### Selection histories and ancient DNA support inheritance through ancestral Native Americans

The genome-wide statistics used above identify candidates with signatures of adaptive introgression but do not directly reconstruct their evolutionary histories. In particular, outlier-based statistics cannot determine whether archaic alleles and alleles showing the strongest positive selection signals occur on the same haplotype, when selection began, or how allele frequencies changed through time. To address these questions, we inferred local ancestral recombination graphs (ARGs), which describe genealogical relationships across the genome while accounting for coalescence and recombination (46). These genealogies allow us to examine which alleles are associated with the selected haplotypes and for estimating their frequency trajectories and selection histories.

We first used the Denisovan-derived *EPAS1* region in Tibetans as a positive control for genealogy-based analyses. Relate (47, 48) recovered strong correlation between Denisovan-derived alleles and alleles with evidence of positive selection within the focal *EPAS1* haplotype (**Fig. 1C**). We then inferred local genealogies for the 25 Peruvian candidate regions using Relate using the combined modern human genome dataset that includes 88 Peruvian genomes and sampled individuals from AFR, EAS, and EUR populations (see methods). When Relate did not provide a usable output, we used SINGER (49) on the 88 Peruvian genome dataset. Across all candidates, archaic-derived alleles were concentrated on the same focal haplotypes carrying the strongest evidence of positive selection, broadly resembling the pattern observed at the *EPAS1* positive control. Additionally, candidate regions all retained deep internal branches that are consistent with an older origin of the archaic haplotypes, while shorter terminal branch lengths within Peruvians were consistent with recent frequency increases in Peruvians that led to more recent coalescence (**Fig. S3**). This concordance supports that positive selection acted on introgressed archaic haplotypes rather than simply occurring nearby, while causal alleles cannot necessarily be distinguished from within a haplotype.

We next used these ARGs to infer changes in selection strength through time. To prioritize SNPs whose allele frequencies are not explained by genome-wide ancestry portions, we used Ohana (50), which models each population’s ancestry components and flags variants whose frequencies deviate from the expectation. U20 alleles above the 95^th^ percentile of the chromosome-wide likelihood-ratio distribution were retained as candidate targets for CLUES2 (**Fig. S4**). Here, we define U20 alleles as derived alleles shared between Peruvians and an archaic reference, absent in the African outgroup (YRI), and present at least 20% frequency in Peruvians. We then used CLUES2 to estimate allele frequency trajectories and selection coefficients across user-defined temporal epochs and varied the breakpoint separating recent from older epochs to identify plausible selection onset windows. We obtained selection onset estimates for 21 of the 25 candidate regions (**Fig 1D, Fig. S5-S8**). In general, older epochs showed less evidence of selection than recent epochs, which is consistent with the expectation that archaic variants persisted before selection increased their frequencies.

We evaluated the accuracy of this Relate-to-CLUES2 framework using simulations with different selection coefficients, onset times, and demographic scenarios, including constant population size, exponential growth, and bottlenecks (**Fig. S9, Table S2**). Under the neutral scenarios, false positives rates remained low. In simulated recent selection scenarios, inferred onset times were generally within 100 generations of the true onset (**Fig. 2A**). Accuracy declined in older selection scenarios, especially when selection started more than 300 generations ago and when selected alleles approached fixation. However, these deep onset times fall outside the temporal window most relevant to adaptive events associated with Native American population history. And we report empirical onset time as windows rather than exact time points.

**Fig. 2.**
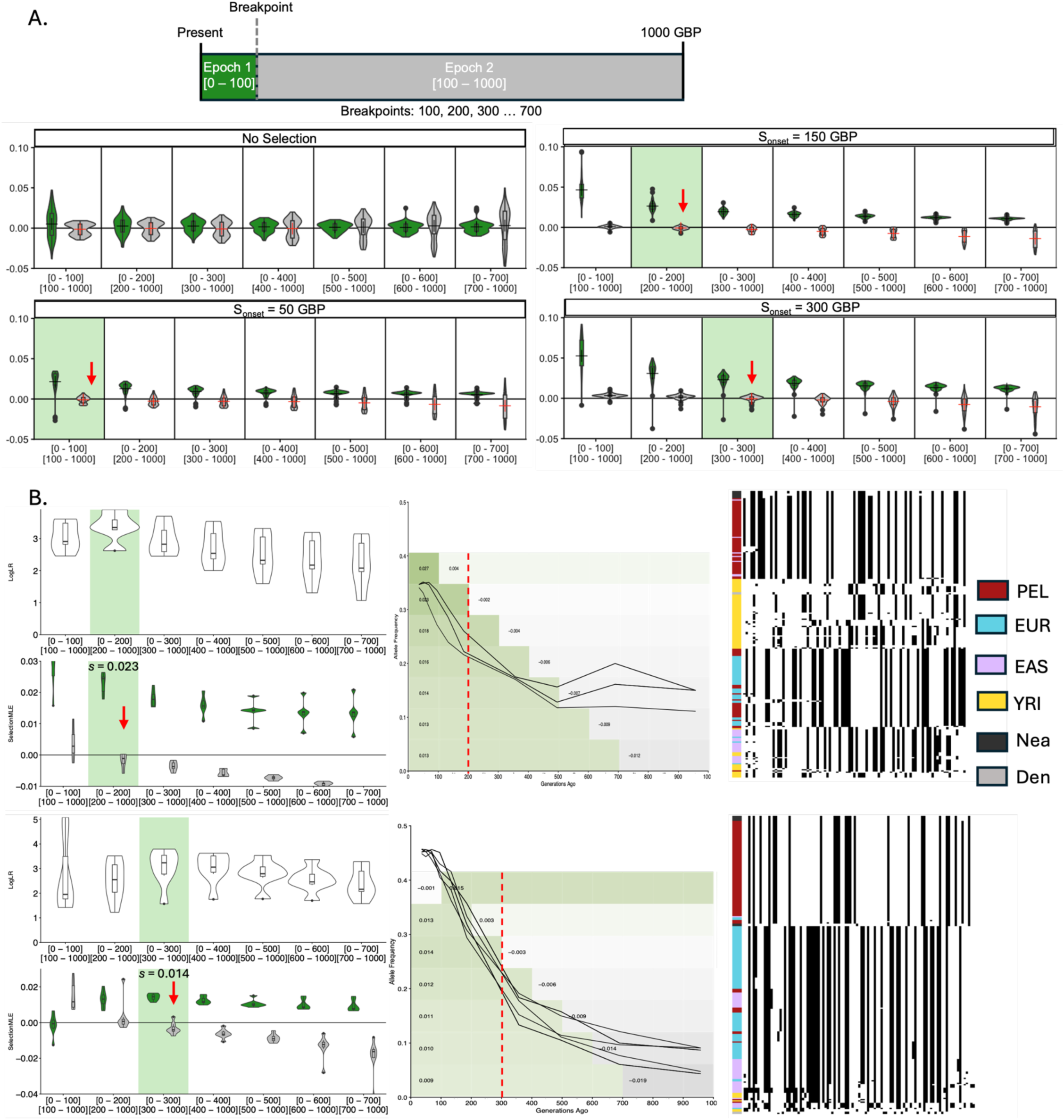
Benchmarking selection onset inference and examples of temporally localized selection. **A)** CLUES2 estimates of selection coefficients across tested epoch breakpoints in simulations. Rows show simulations with no selection or selection onset at 50, 150, or 300 generations before present. Columns represent tested breakpoint models, with epoch 1 corresponding to the recent interval from the present to the tested breakpoint and epoch 2 corresponding to the older interval. Green shading marks the epoch containing the true simulated selection onset, and red arrows indicate the expected breakpoint or transition. Violin plots show the distribution of inferred selection coefficients across replicates. **B**) Representative candidate loci showing evidence of temporally localized selection. For each example locus, the left panels show epoch-specific support for selection across 100-generation intervals from the present to 1000 generations ago. Violin plots show the distribution of log-likelihood ratios (LogLR) comparing selection versus neutrality, and the corresponding maximum-likelihood estimates of the selection coefficient (SelectionMLE). The inferred selected epoch is highlighted in green with the first inferred negative selection coefficient in epoch 2 indicated with a red arrow, and the estimated selection coefficient indicated in the plot (s = 0.023 for the upper locus; s = 0.014 for the lower locus). The middle panels show inferred allele-frequency trajectories through time, with the green shading marking the epoch of strongest selection support. Each column is a polymorphic SNP, each row is a single haplotype and the colored column to the left corresponds to the population identity of each haplotype (Black: Neanderthal, Gray: Denisovan, Purple: EAS (CHB or CHS), Blue: EUR (CEU or GBR), Red: PEL, Yellow: YRI). The dark cells represent the presence of the derived allele and white space represents the ancestral allele.

For all 21 loci with resolved selection histories, the inferred onset of selection predated the arrival of Europeans in the Americas. Additionally, ancient American genomes provided an independent line of evidence for the presence of candidate archaic haplotypes before post-Columbian admixture. For all 25 focal haplotypes, at least a subset of the defining archaic alleles in the haplotypes were observed in one or more pre-Columbian ancient American genome (**Table S3**; 20, 35-39). These observations support the pre-European contact presence of archaic variation at all 25 regions, whereas the ARG-based analyses independently support the pre-European contact selection at the 21 dated regions. Overall, these findings are consistent with a scenario where the archaic haplotypes were inherited through ancestral Native American populations, whose ancestry passed through the ancient Beringia (51, 52), rather than first being introduced to the Americas through European contact.

Interestingly, present-day local ancestry assignments did not always track this deeper history. Although most focal haplotypes showed East Asian-related ancestry, as expected from Asian-related ancestry of the ancestral Native Americans, four candidate regions were assigned to European ancestry despite the presence of archaic alleles in pre-Columbian ancient genomes and evidence that selection began before the European contact (**Fig. S10**). However, because our ChromoPainter reference panel lacks Indigenous American donors, these assignments may reflect shared ancestry between ancestral Native Americans and Western Eurasians rather than recent European admixture (51, 52). At these loci, restricting local ancestry inference to haplotypes that carry the archaic alleles led to an increase in European ancestry assignment (e.g. *NMUR2* and *MATN2*, **Fig. S10B**). This European affinity suggests a potentially deeper shared ancestry between European and Native Americans, and local ancestry assignments based on modern human reference populations therefore do not necessarily identify the evolutionary origin of archaic variation in the Americas.

### Adaptive introgression was temporally structured across Peruvian population history

The 21 resolved selection onset windows spanned across a broad period of Peruvian history (**Fig. 1D**), from 100 generations ago (2,900 years ago assuming 29 years/generation) to 700 generations ago (20,300 years ago). The midpoint of these windows showed two loose temporal concentrations, with an earlier one near the initial settlement of Peru and a more recent one closer to the Agricultural transition. To test the hypothesis that the selection onset times are temporally dynamic, we applied three statistical tests to distinguish non-uniformity from discrete modes. We used the Cramér-von-Mises test and show that the midpoint estimates departed significantly from a uniform distribution over the tested interval (Cramér-von-Mises =0.605, *p*-value = 0.02), and all 1,000 interval-sampling iterations also returned *p*-values < 0.05. We next used the Hartigan’s dip test to look for support for discrete modes, and in this case, we show that although the midpoint estimates returned significant result (D=0.119, *p*-value=0.00916), only 13.2% of interval-sampling iterations remained significant. Similarly, a two-component Gaussian mixture model provided a modest improvement over a one-component mixture (ΔBIC = 2.388), with ΔBIC>2 in only 5.3% of iterations. Thus, adaptive introgression in Peruvians was strongly temporally structured, but the analyses did not resolve a fixed number of discrete waves.

### Adaptive introgression in Peruvians shows functional effects at multiple biological levels

We further asked whether archaic variants associated with adaptive introgression in Peruvians were concentrated in biological pathways. Using the maximum Q95 values in genes as scores in Signet (53), which tests for selection across connected gene networks, we searched the KEGG database (54) and identified five pathways that were significant in at least four of the five replicate searches: long-term depression, arachidonic acid metabolism, glycerophospholipid metabolism, ether lipid metabolism, and linoleic acid metabolism (**Fig. 3A, Table S4**). A complementary Enrichr (55-57) analysis of genes in the top 1% of the genome-wide Q95 scores recovered arachidonic acid metabolism and linoleic acid metabolism, alongside other lipid-, inflammatory-, neuronal- and chromatin-related pathways, including alpha-linolenic acid metabolism, choline metabolism, inflammatory mediator regulation of TRP channels, ATP-dependent chromatin remodeling and serotonergic synapse (**Fig. S11**).

**Fig 3.**
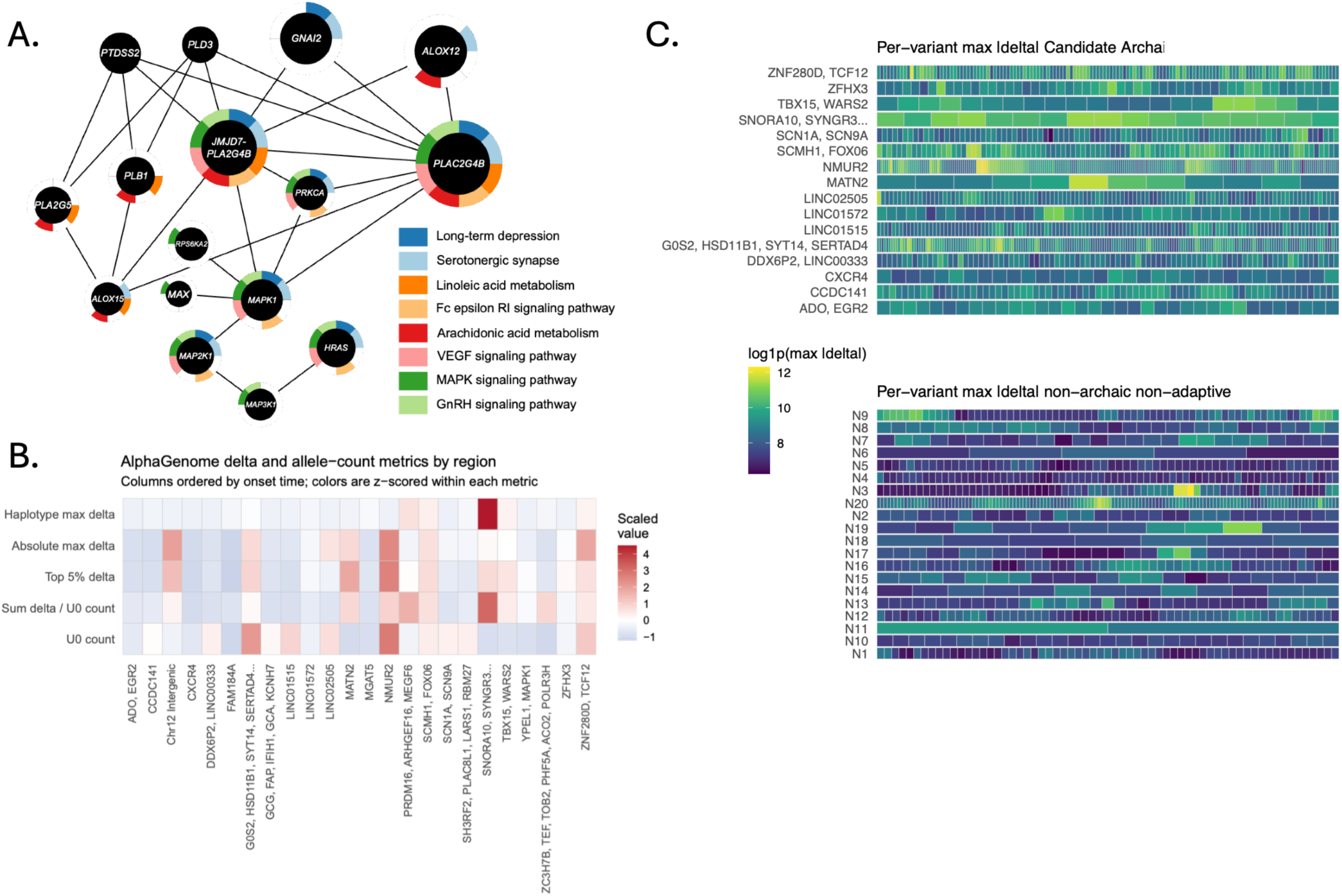
Functional pathways and predicted regulatory consequences of adaptive introgression in Peruvians. **A)** Signet gene-network analysis. Nodes represent genes, colored rings indicate pathway membership, and edges indicate genes with shared network relationships. **B)** Heatmap of AlphaGenome prediction and archaic allele count summaries across candidate regions. Columns represent focal haplotypes ordered by inferred selection onset time when available, and rows represent summary metrics, including haplotype maximum delta, absolute maximum delta, top 5% delta, sum of delta per U0 count, and U0 allele count. Measures are z-scored within each metric, with red indicating higher-than-average values and blue indicating lower-than-average values. **C)** Per-variant AlphaGenome delta summaries in candidate and non-archaic non-adaptive control regions. Stronger and weaker predicted effects are indicated by the plot color scale. Rows represent genomic regions and columns represent individual variants within each region.

Arachidonic acid and linoleic acid metabolism were recovered independently by both methods, providing the clearest convergence on lipid-related pathways (**Fig. 3A, Fig. S11**). Several genes in the candidate regions also connected lipid metabolism with neuronal or inflammatory signaling functions, including *PLA2G4B*, which participates in fatty-acid release from membrane phospholipids (58), and *MAPK1*, which appeared in long-term depression and related signaling pathways. These results suggest that adaptive introgression in Peruvians was not restricted to isolated loci but contributed to sets of genes showing evidence of polygenic selection across multiple biological pathways.

We next examined predicted regulatory consequences of the 25 candidate haplotypes using AlphaGenome (59). Overall, predicted effects were heterogenous, and several candidate regions contained variants with stronger predicted regulatory perturbation scores than non-archaic, non-adaptive control regions (**Fig. 3B-C**). Five candidate regions exceeded the control-derived 95^th^-percentile threshold for the maximum absolute predicted delta, including *ZNF280D*/*TCF12*, *TBX15*/*WARS2*, *SCMH1*/*FOXO6*, *PRDM16*/*ARHGEF16*/*MEGF6*, and *SNORA10*/*SYNGR3* (**Fig. S12**). In several regions, large, predicted effects were concentrated in relatively few archaic variants rather than scaling with the total number of archaic variants within a haplotype. These computational predictions suggest that regulatory variation represents another potential functional consequence of adaptive introgression in Peruvians, although the predictions do not establish regulatory effects *in vivo* or identify causal targets of selection.

### CXCR4 is a recent immune-related candidate of adaptive introgression carrying a missense variant

Among the candidate regions, the *CXCR4* region emerged as a notable example of recent selection. The haplotypes in Peruvians form a distinct archaic-like haplotype cluster that is not seen in other modern human populations (**Fig. 4A**), and genealogy-based analyses inferred a selection onset at 200 generations ago (approximately 5,800 years ago; **Fig. 4B**), within the more recent concentration of selection onset estimates.

**Fig. 4.**
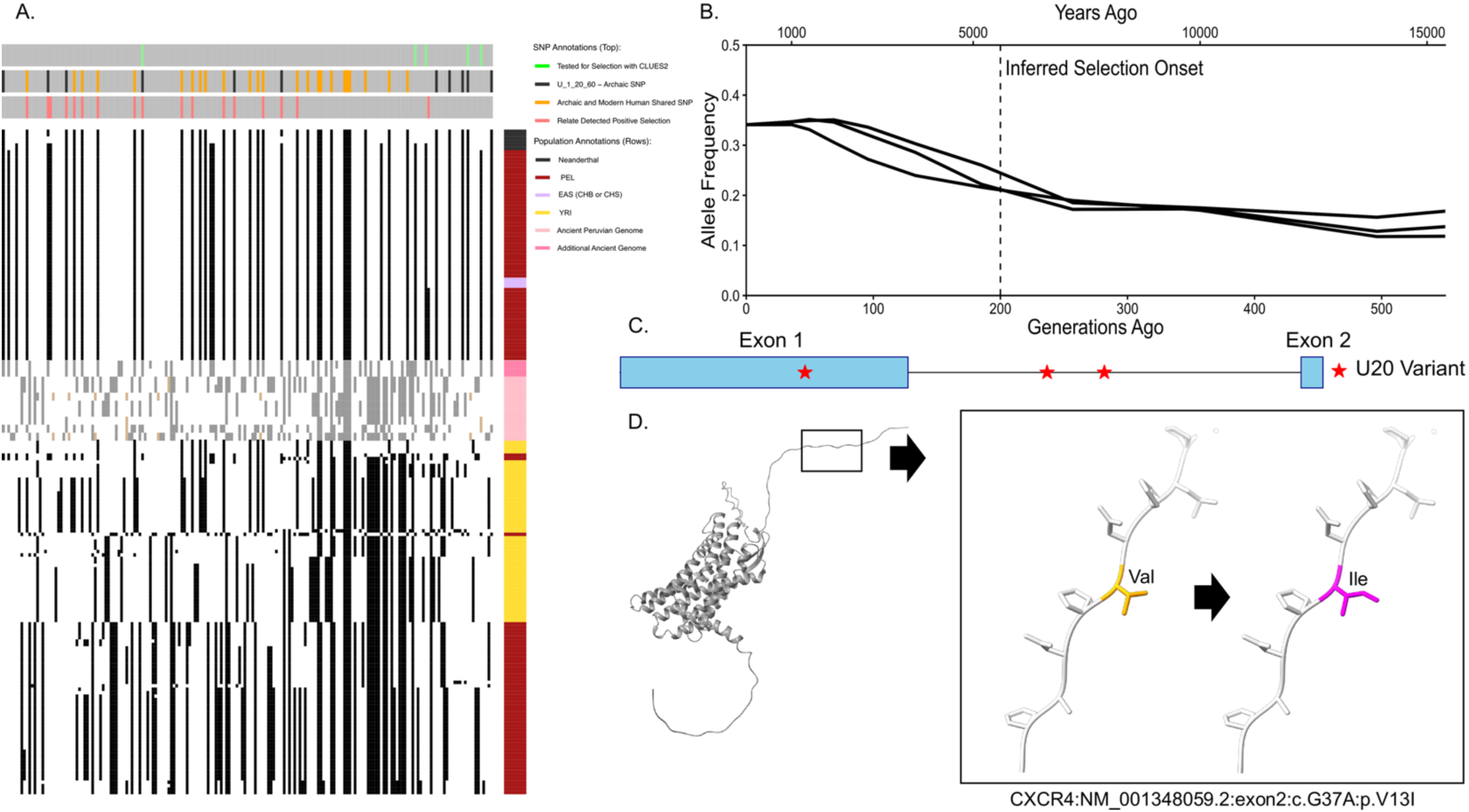
CXCR4 illustrates a case of recent adaptive introgression with coding variation. **A)** Haplotype structure across the candidate region. Rows represent sampled haplotypes and columns represent segregating variants. Black blocks indicate alternate alleles. Colored annotation indicates sampled modern human populations or archaic references. **B)** Relate-inferred allele frequency trajectories and CLUES-inferred selection onset window. The x-axis shows time in generations ago, with approximate years ago shown on the upper axis. **C)** Gene model of CXCR4 showing archaic-associated variants including the V13I coding substitution. Blue boxes indicate exons and red stars indicate candidate archaic variants. **D)** Predicted structural context of the V13I substitution near the N terminus of CXCR4. The inset shows the local structural environment of the substitution, where the non-archaic valine residue is replaced by the archaic-associated isoleucine residue.

The focal haplotype carries a nonsynonymous V13I (**Fig. 4C, D**) substitution near the extracellular N terminus of CXCR4, a domain involved in ligand recognition and receptor activation. *CXCR4* is involved in immune-cell trafficking and inflammatory signaling and is also a major co-receptor for HIV cellular entry (60, 61). However, our data cannot identify HIV or other specific infectious agent as the selection pressure, nor can they distinguish V13I from linked variants as a causal target. Nevertheless, this locus provides a biologically interesting candidate for experimental tests of the molecular and phenotypic effects of selected archaic haplotypes, as the combination of recent selection, archaic-like haplotype structure, immune and environmental relevance, and potentially functional coding variation makes *CXCR4* particularly interesting for functional investigations.

## Discussion

In this study, we refined the landscape of adaptive introgression in modern Peruvians to 25 high-confidence candidate regions and reconstructed selection histories for 21 of them. Three major conclusions emerge. First, all archaic haplotypes were present in the Americas before European contact, and the 21 resolved selection onset times also predate the post-Columbian admixture, supporting their inheritance through ancestral Native American populations. This was likely via the Beringian populations that carried Neanderthal and Denisovan ancestry. Second, selection on different archaic haplotypes acted at different times in Peruvian population history, showing that introgressed variation could remain available long after the original introgression events. Third, functional effects manifest at several levels, including polygenic selection across lipid, inflammatory, immune-related, and neuronal pathways, elevated predicted regulatory effects across archaic haplotypes, and occasional coding variants with potentially adaptive biological consequences. These results support a model in which introgressed archaic variation served as a long-term reservoir that was repeatedly recruited by selection as populations moved into and adapted to new environments in the Americas.

Our results further suggest that adaptive introgression in Peruvians was temporally structured rather than concentrated around a single period of Peruvian history. Although our statistical analyses did not strongly support for a fixed number of discrete modes, midpoints of the inferred selection onset windows loosely separate into two concentrations: an earlier group overlapping the initial settlement of Peru and a later group closer to the transition toward agriculture. We do not interpret these are two clearly separated waves of selection. However, the weak support from the dip test and Gaussian mixture modeling likely reflects the limited number of resolved loci. Nevertheless, the observed distribution was significantly non-uniform, and their apparent overlap with major ecological and cultural transitions may not be coincidental. Such transitions substantially changed the environment and lifestyles experienced by Peruvian populations (24), potentially altering the fitness effects of archaic variants that remained as neutral standing variation.

The earlier concentration overlapped with the settlement of the Andean region, during which populations encountered high-altitude hypoxia (28), dietary shifts to reliance on fishing resources along the Peruvian coasts (30), and increasing use of locally available vegetation including tubers and maize (29, 31, 32). The later concentration, in contrast, occurred closer to the transition toward agriculture and more stable lifestyle, accompanied by changes in population size, social structure, and exposure to infectious disease. Any of these shifts could have changed the fitness effect of archaic variants already segregating in the population. Even if we cannot assign individual selection event to specific ecological or cultural changes, these historical contexts provide plausible historical contexts for the different onset times.

*CXCR4* provides a particularly interesting example from the more recent period. Its archaic-like haplotype has an inferred selection onset around 200 generations ago when agriculture started and carries the V13I missense substitution near the extracellular N terminus. CXCR4 has broad roles in immune-cell signaling and is also well known as a major HIV co-receptor (60, 61). There is no established link between the selection in Peruvians and HIV, but the relatively recent selection timing raises the possibility that this locus responded to the changes in infectious disease pressures associated with agriculture and population growth, which favored a change in immune function. We do not know the specific pathogen or phenotype involved, or whether selection acted on V13I or linked variation. However, the selection could have involved a viral or infectious agent using pathways that overlap with those later exploited by HIV. These uncertainties make this locus a highly interesting target for future functional experiments to reveal the molecular mechanism.

Pathway-level findings provide a broader functional context. Signet and Enrichr recovered overlapping signals on lipid metabolism and inflammatory pathways, together with neuronal and other signaling functions. These pathways are particularly interesting in the context of Peruvian population history because lipid biology connects several of the ecological pressures discussed above, including diet, inflammation and hypoxia. Variation in the arachidonic acid pathway has previously been associated with dietary adaptation in Indigenous American populations (16) and Siberian populations (62), both of which historically relied on fat-rich animal resources. More recently, comparing athletes from Andean highland and lowland have identified differences in linoleic acid and arachidonic acid metabolism, including differences in free fatty acids and downstream metabolites suggesting these pathways contributing to altitude-related physiological responses (63). The pathway tests were genome-wide, and they should not be interpreted as functional validation of the 25 focal regions.

Regulatory predictions provide another level of functional information. We show that some archaic haplotypes had larger predicted AlphaGenome effects, often driven by a small subset of variants. This suggests that the functional consequences of adaptive introgression were not limited to coding variations and may frequently have involved changes in gene regulation. At the same time, occasional missense variants such as the V13I substitution in *CXCR4* provide examples in which adaptive introgression directly affect protein structure. The functional effects and phenotypes under selection all remain to be tested experimentally.

Another observation is that present-day local ancestry does not always reflect the deeper origin of archaic haplotypes. Specifically in our study, four of the 25 candidate loci showed greater affinity to European reference haplotypes despite evidence that the archaic alleles were present in the Americas and under selection before European contact. Interestingly, archaic-like haplotypes at all 25 loci were also found in European and/or East Asian populations, generally at lower frequencies than in Peruvians (Table S1). Although some loci showed suggestive evidence of selection in these populations, none passed the significance thresholds used in our analyses. This suggests that archaic variation was shared among Eurasian and ancestral Native Americans but may have experienced different selection histories. The European ancestry assignments could reflect deeper shared ancestry, limitations in our reference panel, or potentially the reintroduction of similar archaic haplotypes through recent European admixture. Thus, local ancestry alone cannot resolve the evolutionary origin of archaic adaptive haplotypes, highlighting the value of integrating ancient DNA and reconstructed selection histories.

Several methodological limitations should be considered when interpreting these results. First, we were only able to resolve selection onset time for 21 of 25 candidate regions using CLUES2. Our simulations indicate that the Relate/CLUES2 framework has low false positive rate under the demographic scenarios relevant for American populations and can recover relatively recent selection onset with useful accuracy, but performance declines for older onsets (e.g. greater than 300 generations ago) and when selected alleles approached fixation. This time range is older than most of the events relevant to Peruvian population history, but it indicates that older inferred selection times should be treated with greater caution. Additionally, for 12 loci, the breakpoint between the strongest likelihood support did not coincide with the transition between epochs inferred to have positive selection and non-positive selection. We represented the difference as an uncertainty interval rather than assigning a single onset date. These cases illustrate that the temporal resolution of CLUES2 and the breakpoint procedure is limited. Future improved inference methods and high-quality time-series data should help resolve selection histories more precisely.

The small number of resolved loci also limits how much we can conclude about the detailed structure of adaptive introgression in the Americas through time. With only 21 onset estimates, we have considerably stronger evidence that selection was non-uniform than evidence for two or other fixed number of waves. Moreover, the statistical null does not itself account for potentially uneven inference power over time or the ascertainment of candidate loci. Additional well-supported adaptive introgression loci, ideally from other Indigenous American populations and more ancient genomes will be needed to determine whether the apparent temporal concentrations observed in Peru occurred elsewhere. Similarly, direct functional validation will be required to identify the target phenotypes and the underlying molecular mechanisms for adaptive introgression.

More broadly, our study highlights that adaptive introgression should be better understood as a temporally dynamic process rather than only a genomic signature. Archaic variation can persist long after introgression, move with populations across large geographical distances, and still become beneficial under environments substantially different from those near the original contact zone. This makes American populations particularly informative for studying the long-term evolutionary fate of archaic introgressed variation. Expanding this framework across indigenous American populations should help determine how general these patterns we observed in Peruvians are and contribute independent temporal information to demographic models that remain difficult to infer using current methods.

## Materials and Methods

### Modern Human Genomes

Modern human genotypes were obtained from the 1,000 Genomes Project Phase III (34) and from the Simons Genome Diversity Project (64). Autosomal VCF files from modern humans were merged using BCFtools v1.13 and subsequently filtered using a bed file mask generated by combining the strict Thousand Genomes Project mask (https://ftp.1000genomes.ebi.ac.uk/vol1/ftp/release/20130502/supporting/accessible_genome_masks/) and the SGDP genome mask (https://reichdata.hms.harvard.edu/pub/datasets/sgdp/filters/all_samples/). We analyzed modern Peruvians in Lima, Peru (PEL), the Yoruba in Ibadan, Nigeria (YRI), the Han Chinese in Beijing, China (CHB) and Han Chinese South (CHS), Utah residents with Northern and Western European ancestry (CEU) and British in England and Scotland (GBR) as reference populations that could contribute genetic material to modern Peruvians. In total, the combined dataset includes 88 PEL individuals, and 100 sampled individuals per above modern human populations.

### Ancient and Archaic Human Genomes

We downloaded three high-coverage Neanderthal genomes (1, 65, 66) and their associated mask files from http://ftp.eva.mpg.de/neandertal/ and the high-coverage Denisovan genome (2) from http://cdna.eva.mpg.de/neandertal/Vindija/VCF/Denisova/. Whole genome fasta files were merged using BCFtools v1.13 (67) and filtered using a combined genome mask.

We included in our analyses ancient American genomes with at least 2x coverage from published datasets **(Table S3**), including four individuals from Peru (35) as well as additional ancient American genomes from Chile (38) and Central Mexico (20). We aligned raw sequence reads to the hg19 reference genome using BWA-MEM (68) and sorted the resulting alignments using Samtools v.1.21 (67). Variants were then called for each genome using GATK v.4.3 HaplotypeCaller (69, 70) with a minimum base quality score of 20. We then combined ancient genomes into a single VCF for downstream analysis with Haplostrips.

### Identification of adaptive introgression candidates

We compiled over 150 putative adaptive introgression candidates in Peruvians (PEL in 1000 Genomes Project) from previous studies of adaptive introgression (8, 13-15, 40). In order to refine putative adaptive introgression candidates, we initially evaluated this broad set of candidates using the MaLAdapt assigned score (0-1) (14). Recognizing that MaLAdapt was trained on Eurasian population history which could introduce additional false positives when applied to American populations, we combined MaLAdapt with empirical statistical measures of adaptive introgression. Specifically, we prioritized genomic windows with a MaLAdapt score exceeding 0.80 in Peruvians and not in European or East Asian populations and required mean sequence divergence ratio (15) below the genome-wide 5^th^ percentile, and Q95 (15) score above the genome-wide upper 95th percentile. We excluded regions with one or fewer segregating sites and compared our identified candidates with previously reported candidates to include additional windows with other evidence of adaptive introgression and obtained a set of 90 candidates (8, 13, 15).

### Haplotype visualization

We visualized haplotype structure of candidate windows using Haplostrips (71) separately using the Denisovan and Neanderthals as reference archaic population, alongside PEL, YRI, CHB, CHS, CEU, GBR samples. Alleles were polarized using the ancestral-state option. Sites not represented in the archaic reference were excluded. Missing genotypes in modern samples were displayed according to the software settings.

For each MaLAdapt candidate, we examined the focal window and expanded the window up-and downstream when necessary to capture the extent of the archaic haplotype. Candidate regions were retained if the Peruvian populations showed a contiguous haplotype cluster with high similarity to the archaic reference that was rare or absent in YRI haplotypes. This step reduced the number of candidate windows from 90 to 40.

We also added ancient American genomes to the haplotype comparisons to assess the presence of defining archaic alleles before the European contact. Ancient genomes were not phased. Each ancient individual was therefore displayed as a single pseudohaploid row rather than two haplotypes, and rows should be interpreted as a random sample of one allele per site, not as true haplotypes. We therefore treat the ancient genomes as evidence for the presence of individual archaic-defining alleles, not for intact haplotypes.

### Selection Scan in Modern Peruvians

We examined evidence of recent positive selection, especially soft sweep signatures, using haplotype-based statistics calculated in Selscan 2.0 (41), including the Number of Segregating Sites by Length (nSL; 43) and the haplotype homozygosity-based method, H12 (44). Additionally, we used the cross-population XP-nSL statistic to specifically test for positive selection in the PEL population when compared to East Asian subpopulations. VCF files were converted to the HAPS/SAMPLE format using PLINK 2.0 (72), and genetic map files were constructed using a modified HapMap with a custom Perl script. We ran Selscan separately for each chromosome, and then the resulting test statistics were normalized across the entire genome for each statistic independently. We defined a significance threshold at the 95th percentile of the normalized distribution for each test. Finally, a genomic region was retained as a candidate if it contained at least one U20 SNP, defined here as derived alleles shared between Peruvians and an archaic reference, absent in the African outgroup (YRI), and present at least 20% frequency in Peruvians, and reached genome-wide significance value (above the most extreme 5th percentile) for other statistics.

### Control regions: non-introgressed genomic intervals

For comparisons, we first filter for genomic regions with low evidence of archaic introgression. These regions are defined by an average maladapt score below 0.7 and an average sequence divergence ratio above the 80% quantile of the genome-wide distribution. Using this set, we assembled two different sets of negative controls, including one set with 20 regions that contain selection signatures and another set with 20 regions without evidence of selection based on Selscan and Relate outputs. We used set of 20 regions for each control to match with the test set size.

### Local Ancestry Inference

We estimated local ancestry for each candidate genome region using Chromopainter v2 (73) which estimates the probability of ancestry (copy probability) by each donor haplotype for each SNP in each recipient haplotype by determining the first coalescing donor haplotype. Local ancestry of each region was classified based on the population of the haplotypes exhibiting the highest mean donor probability for the Peruvian haplotypes with archaic ancestry. For candidate regions, we compare the full population mean to the mean copy probability of the subset of PEL haplotypes that carry the archaic haplotype for each region. This approach allowed us to pinpoint the specific contribution of donor populations to the archaic introgressed segments in comparison to non-archaic haplotypes. We expected this to represent the ancestral population (EAS, EUR, or AFR) that introduced each haplotype to AMR populations, though in highly admixed populations, like Peruvians, local ancestry turn-over may confound this inference.

When running Chromopainter v2, we assigned the 176 PEL haplotypes as the recipient population. The donor populations were chosen to represent the major ancestral lineages contributing to modern Peruvian ancestry: East Asians, Europeans, and Yoruba. The combined modern human VCF files were converted to ped/map files using PLINK 2.0 and then converted to phase format using the provided perl script plink2chromopainter.pl. Recombination map files were generated using convertrecfile.pl with human genome hapmap files.

We estimated the nuisance parameters, mutation rate (mu) and effective population size (Ne) using the Expectation-Maximization algorithm on all candidate regions. We used the -b switch to produce output with the copy probability of ancestry from the three assigned donor populations (EAS, EUR, and YRI) for each locus for each haplotype. The portion copied across the region for each source population is averaged across all PEL haplotypes for each locus and then plotted across the full length of the archaic haplotype.

### Identifying population-specific selection signatures with Ohana

We used Ohana (50) as a tool to evaluate ancestry-specific signatures of positive selection by first assessing population structure to identify sites whose allele frequencies cannot be explained by ancestry components. We estimated chromosome-wide covariance matrices using K=4 ancestral components and identified focal U20 loci with likelihood ratio scores above the 95^th^ percentile of their chromosome-wide distribution.

### Selection History Inference

We inferred local genealogies and allele frequency trajectories using RELATE v1.2.2 (47, 48), and inferred selection histories using CLUES2 (74, 75). Genotypes were converted to haps/sample format using the RelateFileFormats pipeline. We then used the module PrepareInputFiles with the phase3 hg37 ancestral genome (https://zenodo.org/records/15801307), the hg37 recombination map files, the haps/sample and poplabels file for polarization. We ran RELATE on each chromosome with a default mutation rate of 1.25e-8 and 30,000 as the effective population size of haplotypes. We estimated population size histories and extracted population-specific genealogies for PEL using RelateExtract. We used the DetectSelection.sh script in PEL, EUR, EAS, and YRI to examine the geographical distribution of selection signature.

For each focal region, we selected at least three high-frequency U20 alleles with MaLAdapt or Ohana support for selection history analyses in CLUES2. CLUES2 uses local genealogies and coalescent rates from Relate to assign an estimated selection coefficient to each user-defined epoch. We ran CLUES2 with 500 importance samples per locus, population-specific allele frequencies, and the PEL-specific coalescence file. We converted genealogical inputs using the RelateToCLUES pipeline.

We used a shifting break-point approach to estimate selection onset. For each selected SNP, seven two-epoch models were fitted, with breakpoints at 100-generation intervals from 100 to 700 generations ago and the oldest boundary fixed at 1,000 generations. For each candidate region, we computed the median log-likelihood ratios (LLRs) across tested SNPs and compared the inferred selection coefficients before and after the breakpoint. When the most supported breakpoint did not agree with the transition of LLR between epochs, we represented the disagreement as an onset uncertainty interval in the timeline.

For regions not resolved with RELATE, we inferred genealogies using SINGER (49) and used the SINGERtoCLUES2 pipeline to estimate selection onset under the same breakpoint procedure above.

We evaluated the 21 resolved selection onset estimates for departure from a uniform distribution across 0–700 generations and for evidence of multimodality. For selection onset times represented as an interval, we used the midpoint for the initial test and sampled within the interval to assess sensitivity to uncertain onset times. We assessed departure from a uniform distribution using the Cramér–von Mises (CvM) goodness-of-fit test in R package goftest (76). In 1,000 Monte Carlo iterations, we sampled onset time independently within each locus-specific interval and repeated the test. We recorded the proportion of iterations with *p-value* < 0.05.

We tested multimodality using Hartigan’s dip test in the R package diptest (77), first with interval midpoint and repeated across 1,000 interval-sampling iterations. We also fitted one- and two-component Gaussian mixture models using the mclust package (78). For each set of onset times, ΔBIC was defined as BIC (two-component) – BIC (one-component), with a positive ΔBIC indicating favor for the two-component model. We recorded the proportion of iterations with ΔBIC >2.

### EPAS1 positive control in Tibetans

As a positive control, we reconstructed the genealogy of the Denisovan-derived *EPAS1* region in Tibetans. 38 Tibetans from the dataset analyzed by (22) and originally collected by (79). We combined these data with Han Chinese (CHB and CHS) and Yoruba (YRI) genotypes described above, and restricted genotypes using genome accessibility masks and used biallelic variants.

We converted the merged VCF to haps/sample format and ran Relate v1.2.2 using the same ancestral genome, recombination map, mutation rate and effective population size as in the Peruvian analyses. We extracted the Tibetan-specific genealogies with RelateExtract and ran DetectSelection.sh to identify SNPs with evidence for positive selection. We defined Denisovan-derived alleles as U20 alleles and compared their location and haplotype background with variants detected with evidence of positive selection and visualized their frequency trajectories across the *EPAS1* region.

### Simulation-based benchmarking

We benchmarked accuracy on two aspects of our inference framework using simulations, including 1) the detection of archaic alleles under positive selection and 2) the inference of selection onset time. Simulations were based on demographic scenarios with changes in population size, which was intended to reflect conditions relevant to American populations.

We simulated 2-megabase (Mb) regions under a three-population demographic model adapted from (80) using SLiM v5 (81). In this model, a small archaic population (N = 100) diverged from the ancestral human population (N = 7,310) approximately 6,000 generations before present. A fixed neutral mutation was introduced in the archaic population before introgression. The ancestral human population further split into a modern African and a non-African recipient population (N = 1,861) 2,056 generations ago, and 20 generations later, a pulse of 10% archaic introgression was introduced to the recipient population. The introgressed mutation remained neutral until the specified selection onset time (50-300 generations ago) after which the selection coefficient was set to *s* = 0.001-0.02 (**Fig. S13**). All simulations were conditioned on the target mutation not being lost in the recipient population, and VCFs were sampled at the end of the simulations for Relate and CLUES2.

In simulations with positive selection, true positive rates (TPRs) were defined as the proportion of simulations where DetectSelection reported a significant U20 variant within 250 kb of the target mutation. False positive rates (FPR) were calculated as the proportion of non-overlapping 50 kb windows outside of the 250 kb interval that contained at least one significant U20 variant.

For onset time benchmarking, we applied CLUES2 to five U20 variants within 100 kb of the target mutation and applied the two-epoch breakpoint procedure described above. The significant log likelihood ratio (LLR) was defined using the 95^th^ percentile from neutral simulations. We assessed inferred onset times across demographic scenarios with and without a target population bottleneck.

### Functional Enrichment of Candidate Regions

We used the R Bioconductor Graphite (82) and Signet (53) packages to test for evidence of polygenic selection in subset of connected gene networks. Signet searches for subnetwork using simulated annealing and compares network scores with a null distribution. We used the maximum Q95 value within a gene as score for Signet and ran it 5 times against a null distribution using 20,000 iterations per run. We retained pathways with a p-value below 0.05 in four or five replicate tests.

As a complementary, non-network-based analysis, we used the web-based Enrichr tool (https://maayanlab.cloud/Enrichr/; 55-57) to look for overrepresentation of genes in biological pathways. We queried the KEGG 2026 library (54) and GO libraries including GO Biological Process and GO Molecular Function. Enrichr reports. We retained pathways with Benjamini-Hochberg adjusted *p-values* < 0.05.

We used AlphaGenome (59) to estimate predicted sequence regulatory effects for archaic variants in candidate and all variant effects in non-archaic control regions. For single-variant scores, reference and alternate alleles were evaluated with the model and summarized as the maximum absolute delta score across reported outputs and positions. For haplotype-level scores, archaic variants from the same interval were introduced jointly as alternate sequence and evaluated together. This allowed AlphaGenome to evaluate the combined sequence effect of the haplotype within a single query.

## Acknowledgment

The authors thank Jeffrey Kidd and the Zhang Lab at the University of Michigan for helpful discussions during the development of this project. XZ, SG and AA were supported by NIH grant R35GM154856 (awarded to XZ).

## Data and code availability statement

All genomic data analyzed in this study were obtained from previously published datasets, including the 1000 Genomes Project, Simons Genomic Diversity Project, and publicly available ancient and archaic human genomes, as described in Materials and Methods. The identified adaptive introgression candidate regions, inferred selection histories and supporting summary statistics are provided in the Supplementary Information and Supplementary Tables. Custom Scripts used for all statistical analyses, inference and simulations are available at https://github.com/xzhang-popgen/PeruvianAI

