## Supplementary material for "Temporally Dynamic Waves of Selection on Archaic Variation in Peruvians": SI Figures

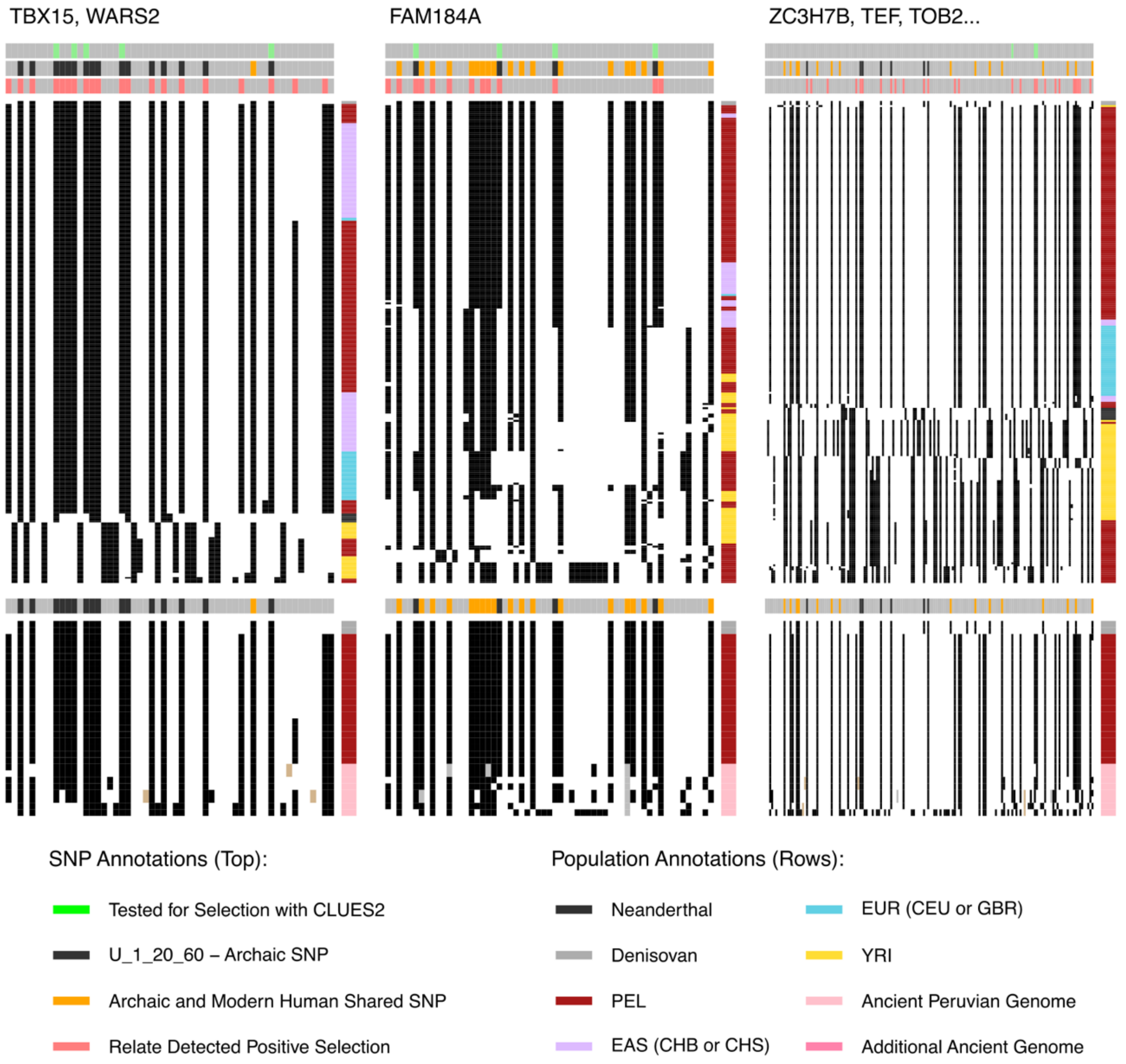


**Fig. S1. Haplostrips illustrating haplotype structure for each candidate archaic locus discussed in this paper.** Each column is a polymorphic SNP, each row is a single haplotype and the colored column to the left corresponds to the population identity of each haplotype (Black: Neanderthal, Gray: Denisovan, Purple: EAS (CHB or CHS), Blue: EUR (CEU or GBR), Red: PEL, Yellow: YRI, Pink: Ancient American genome). The dark cells represent the presence of the derived allele and white space represents the ancestral allele.

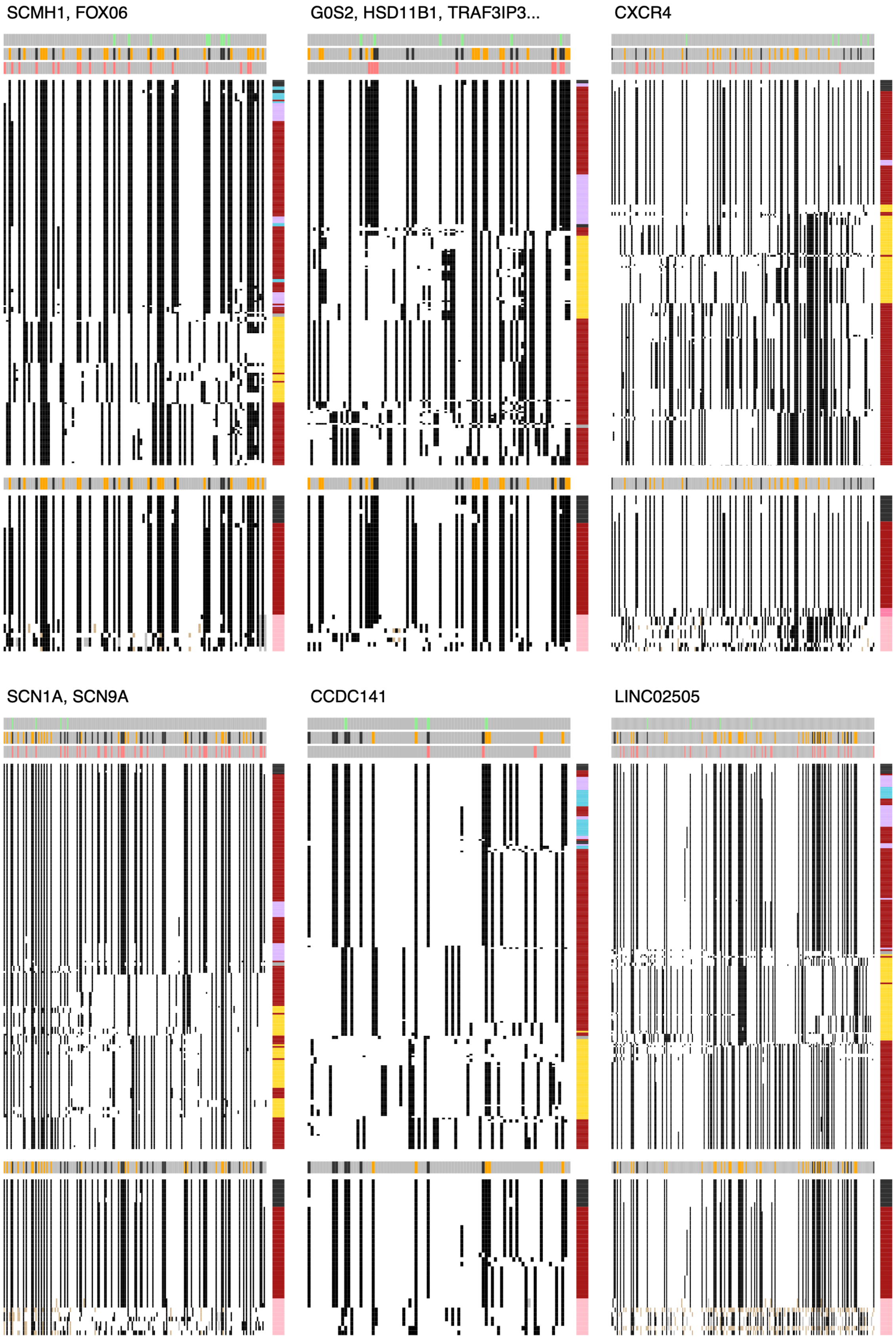


**Fig. S1.** (Continued)

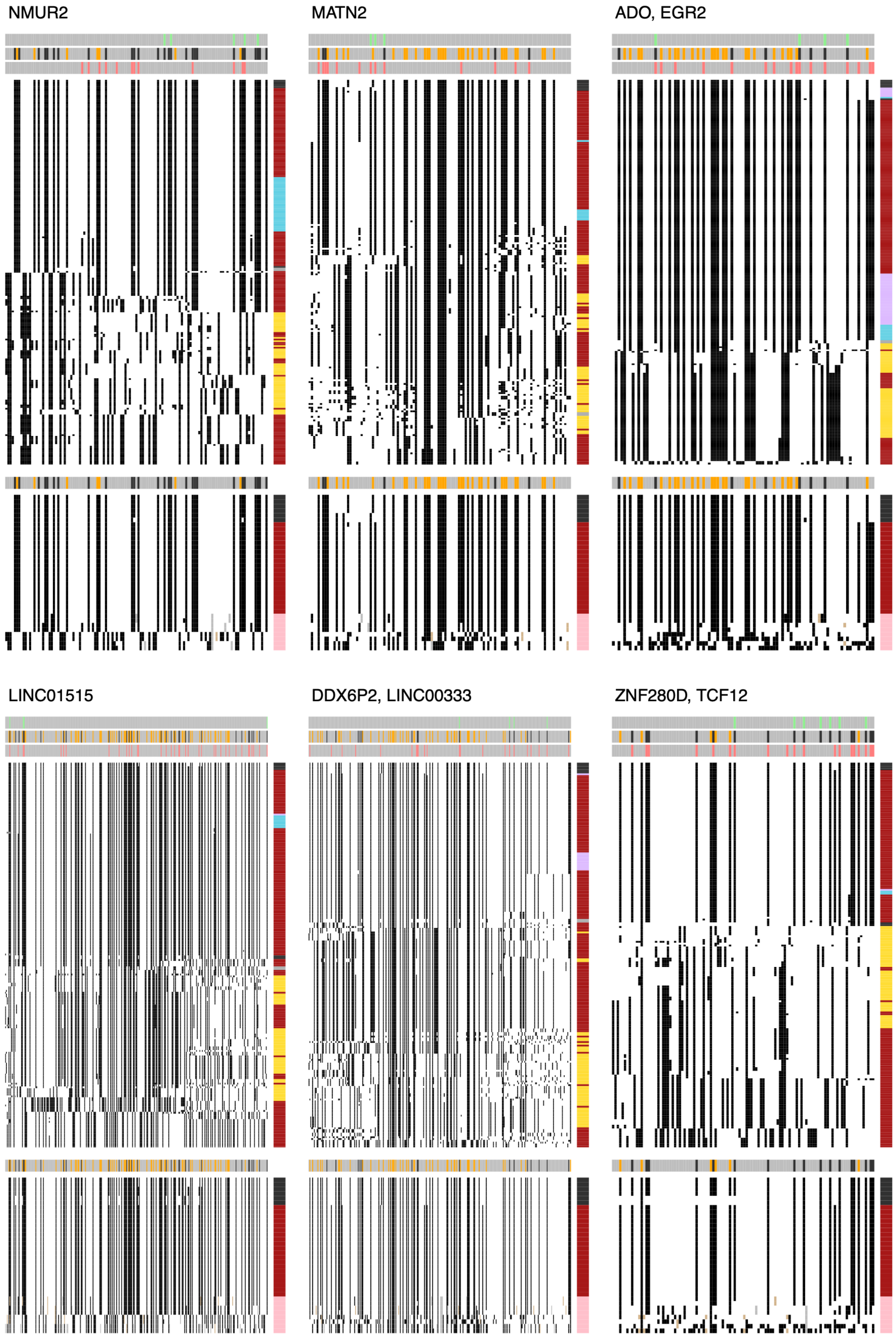


**Fig. S1.** (Continued)

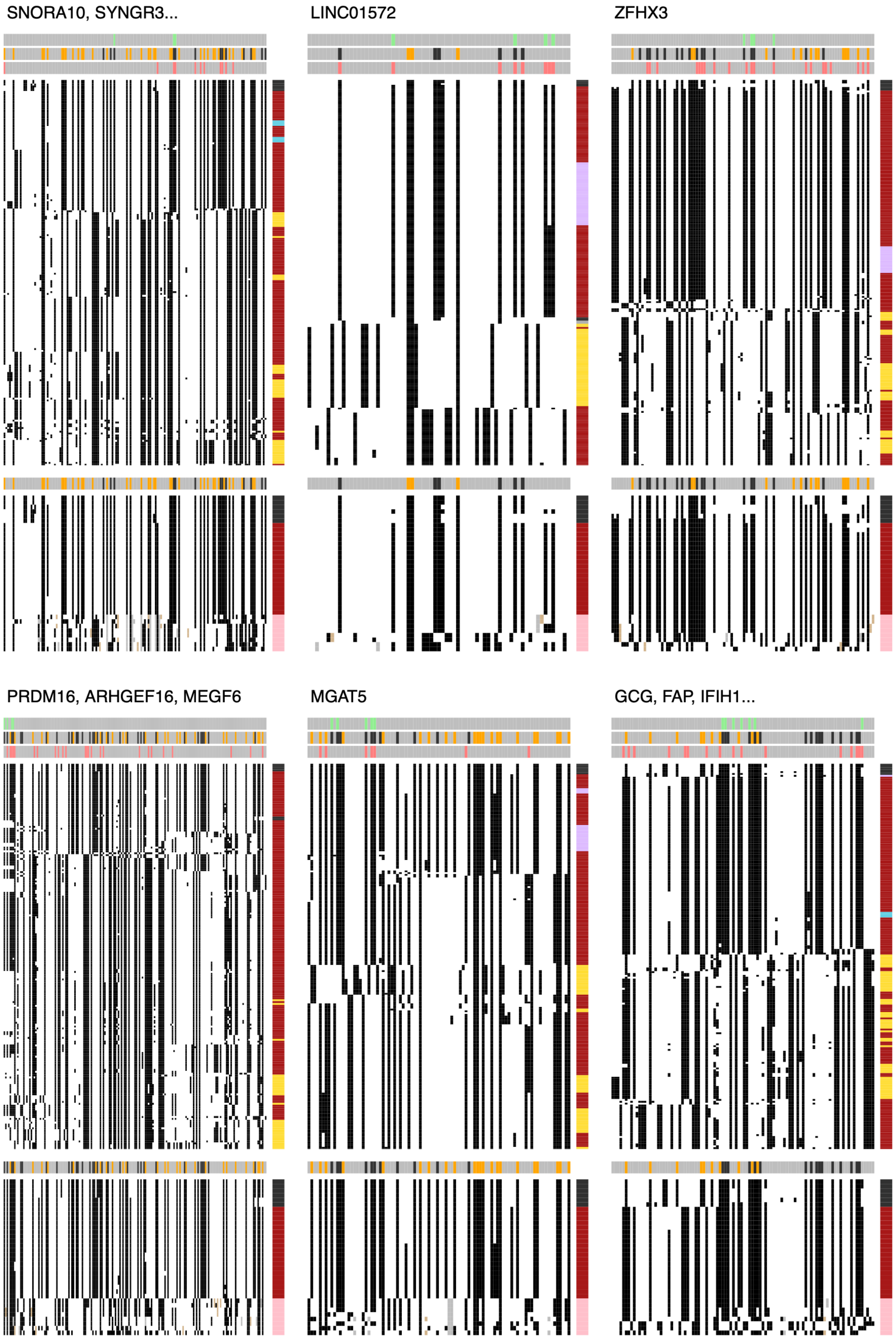


**Fig. S1.** (Continued)

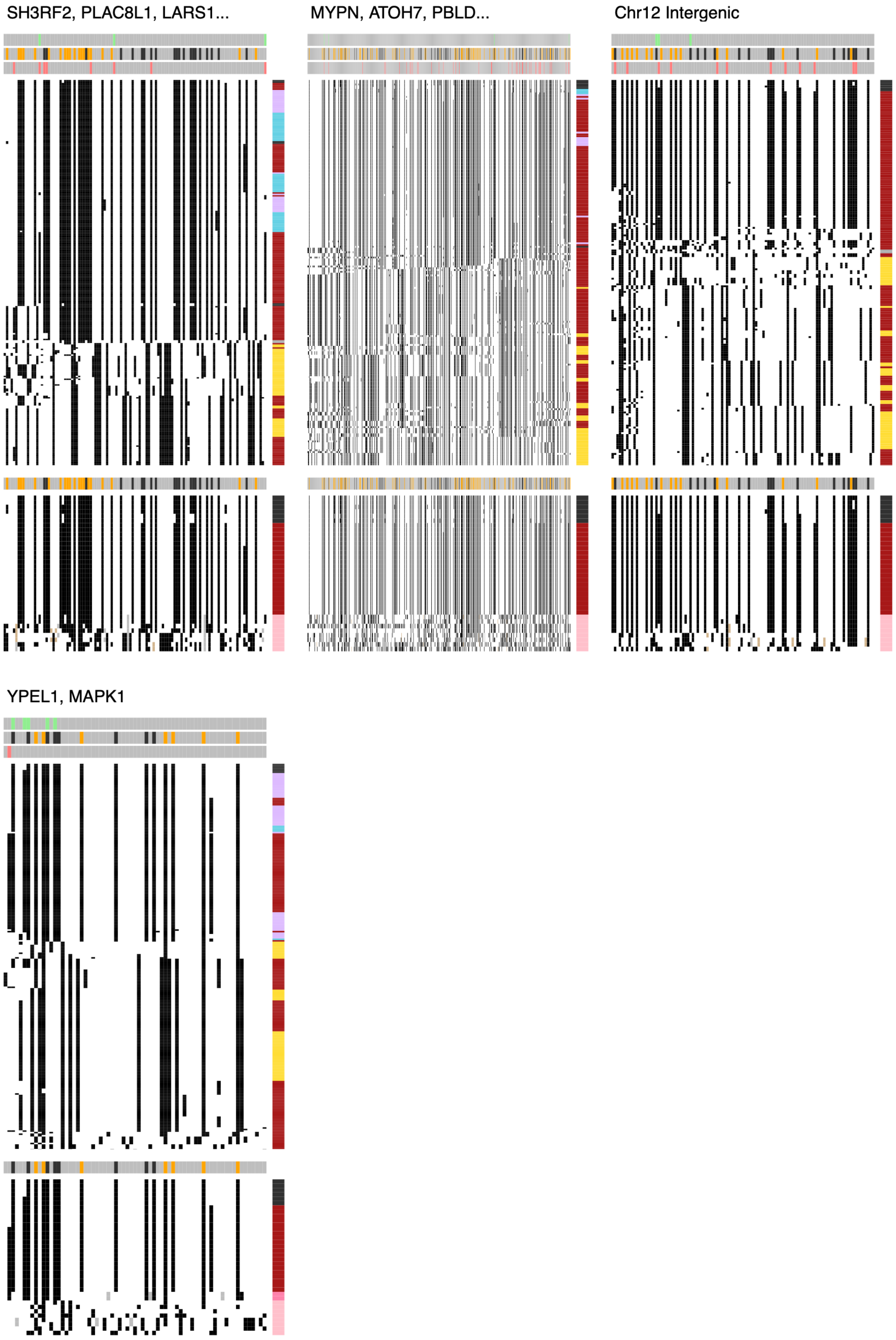


**Fig. S1.** (Continued)

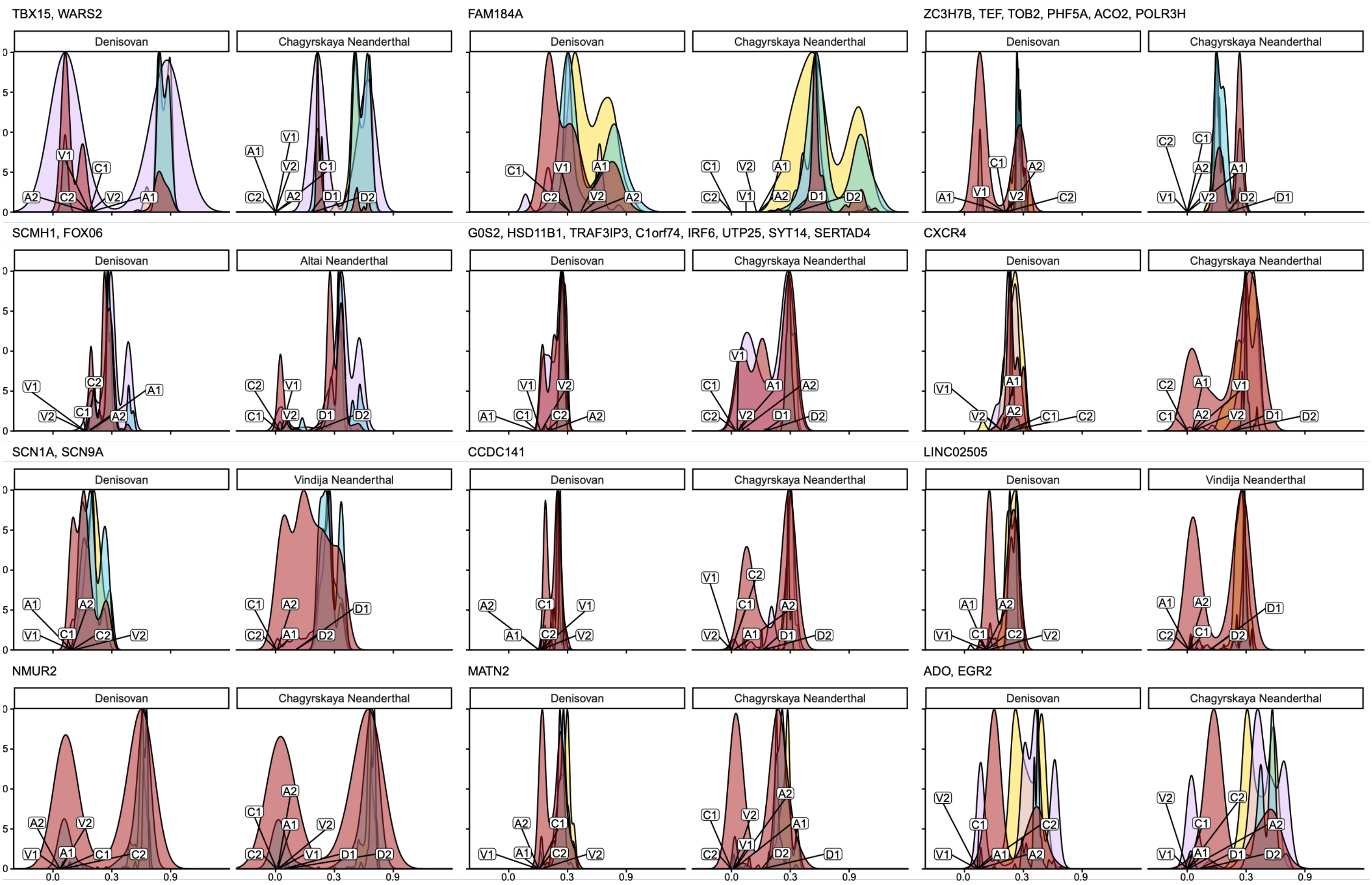


**Fig. S2. Genetic distance between modern haplotypes and archaic genomes**. Separate plots are shown for the Denisovan genome and the single Neanderthal genome that shows the lowest divergence to modern Peruvian haplotypes. The distance between the reference genome and each archaic haplotype is also plotted. Colors correspond to population groups as follows: Red – PEL; Yellow – YRI; Purple – EAS (CHB and CHS); Blue – EUR (CEU and GBR).

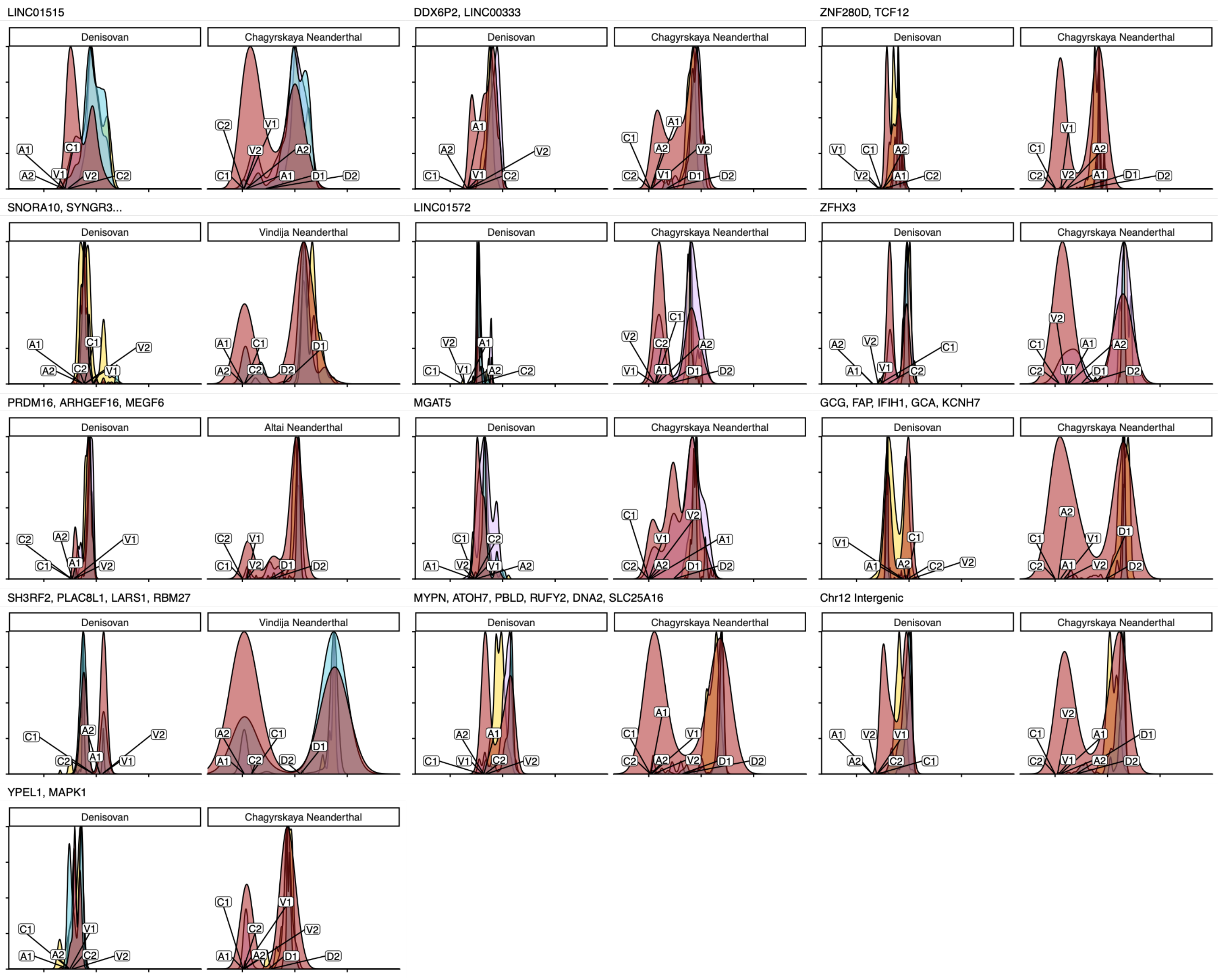


**Fig. S2.** (Continued)

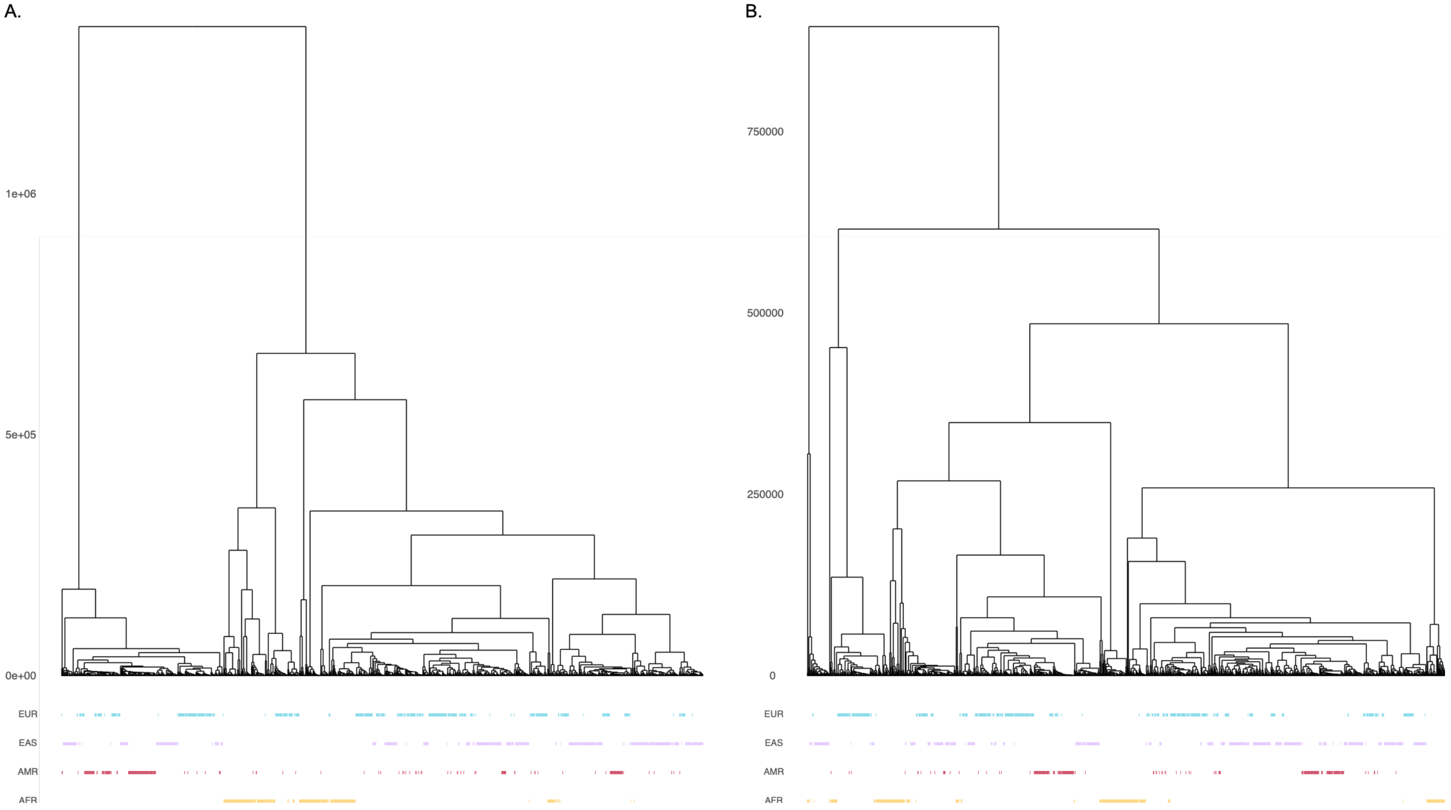


**Fig. S3. Relate-inferred genealogies illustrating differences between archaic SNPs under selection in Peruvians to non-archaic SNPs with no evidence for positive selection in Peruvians**. A) Local genealogy at Chr5:145,474,584 (hg19) within the *SH3RF2*/*PLAC8L1*/*LARS1*/*RBM27* candidate region, an archaic-like SNP with evidence of positive selection in Peruvians. B) Local genealogy at Chr5:64,661,169 (hg19), a non-archaic SNP located within an intron of ADAMTS6 with no evidence for selection within Peruvians, shown as a control. The y-axis shows coalescene time in years before present (assuming 28 years/generation). Tips represent individual haplotypes and colored bars beneath each tip mark the population of each modern haplotype: Red – PEL; Yellow – YRI; Purple – EAS (CHB and CHS); Blue – EUR (CEU and GBR). Peruvian haplotypes carrying the archaic-like alleles cluster together on a deep internal branch with short terminal branches, consistent with an older origin and a recent frequency increase in Peruvians, and they are absent or rare among AFR haplotypes.

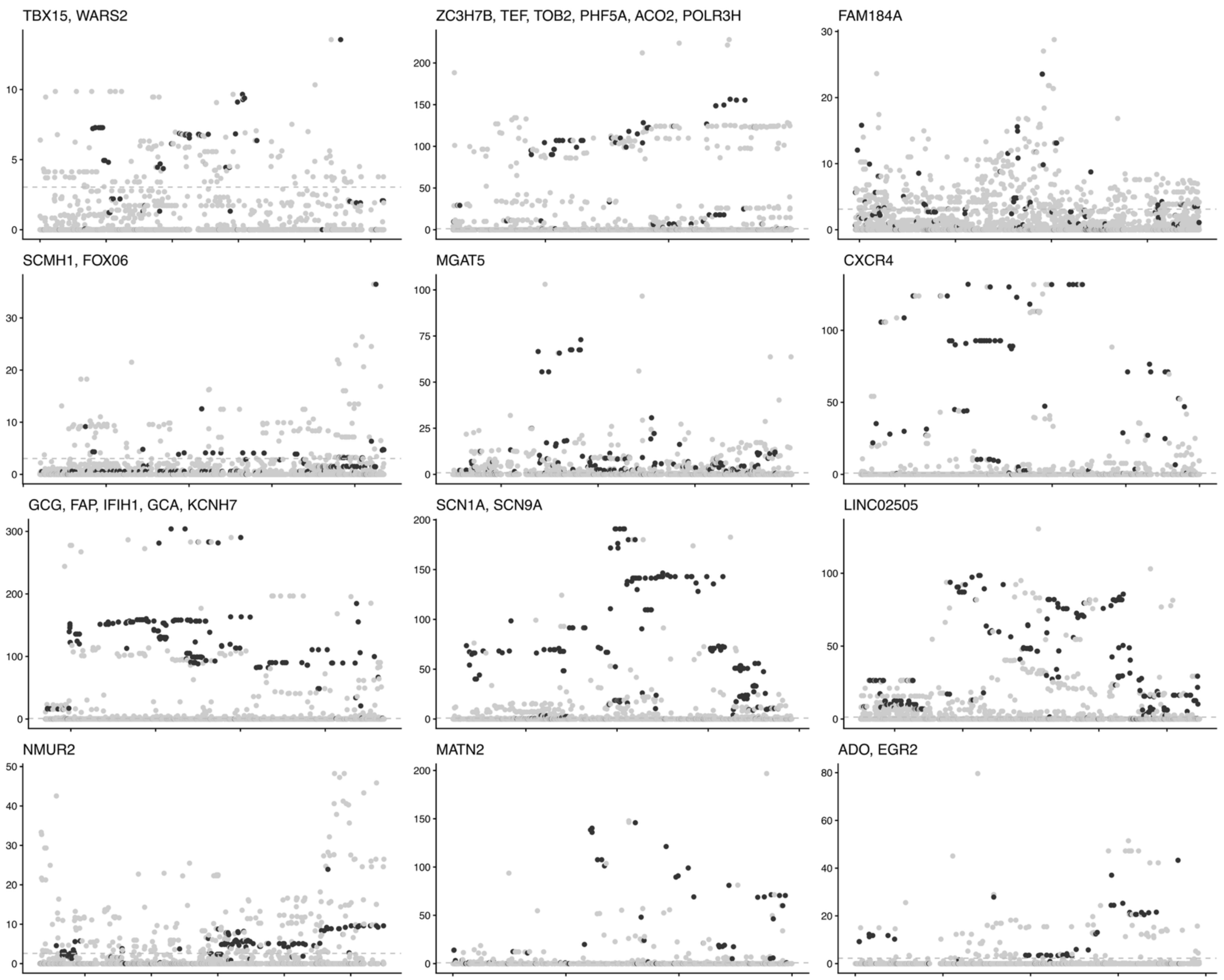


**Fig. S4. Ancestry-aware selection scan with Ohana across candidate archaic haplotypes in Peruvians.** Ohana selection scores for individual SNPs within each focal archaic haplotype. Each panel shows one candidate region, with genomic position on the x-axis (hg19) and the Ohana likelihood-ratio (LLR) score on the y-axis. Each point is a SNP within the haplotype. Black points represent U20 loci, and all other SNPs are plotted in gray. Scores were computed from allele frequencies in PEL, modeled with K = 4 ancestral components. This tests whether a SNP's frequency departs from what the population's ancestry structure alone predicts. The gray dashed line marks the 95th percentile of the chromosome-wide LLR distribution for the corresponding chromosome. SNPs above the line are outliers for ancestry-specific selection. U20 alleles that exceeded this threshold were prioritized for selection-history inference with CLUES2.

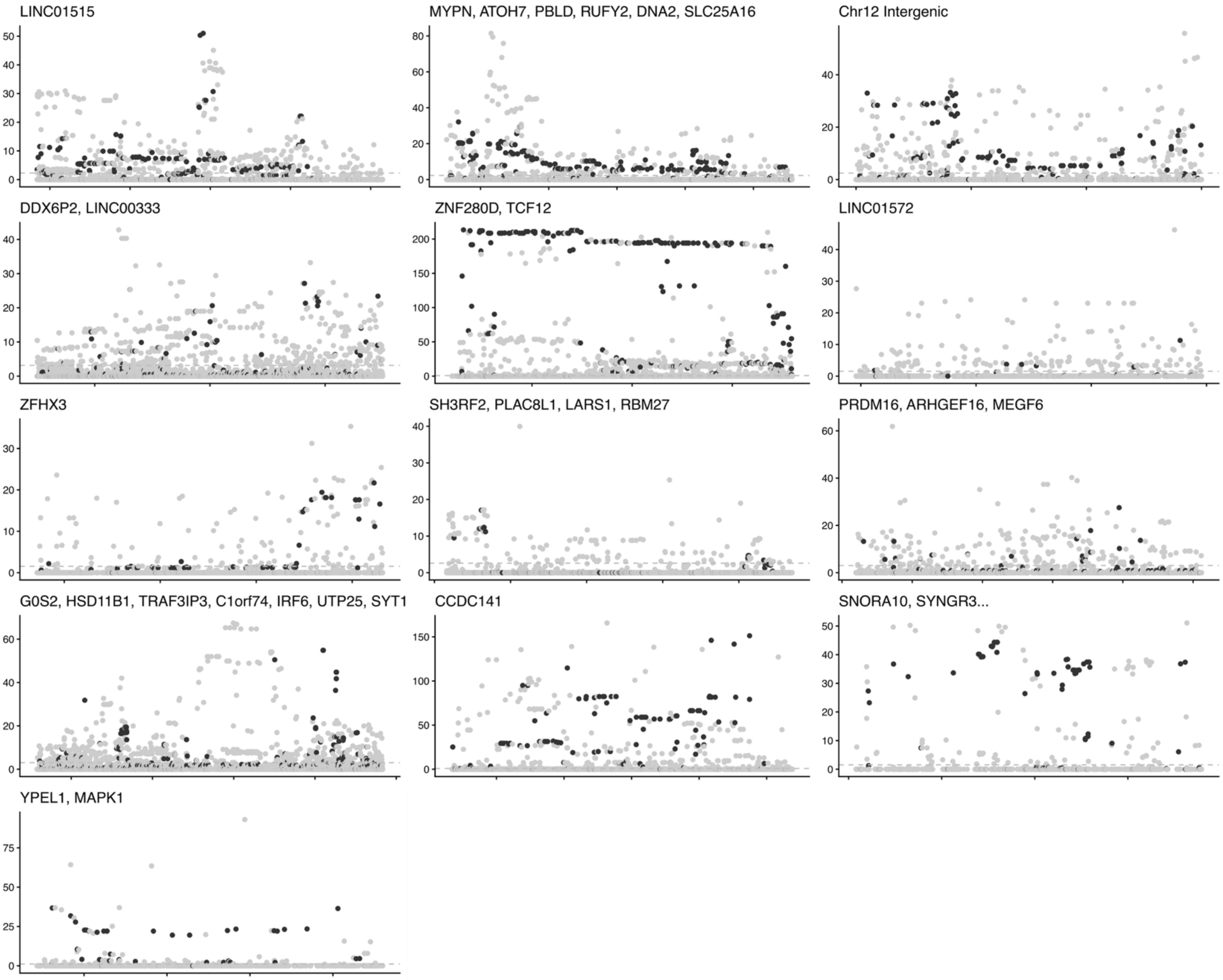


**Fig. S4.** (Continued)

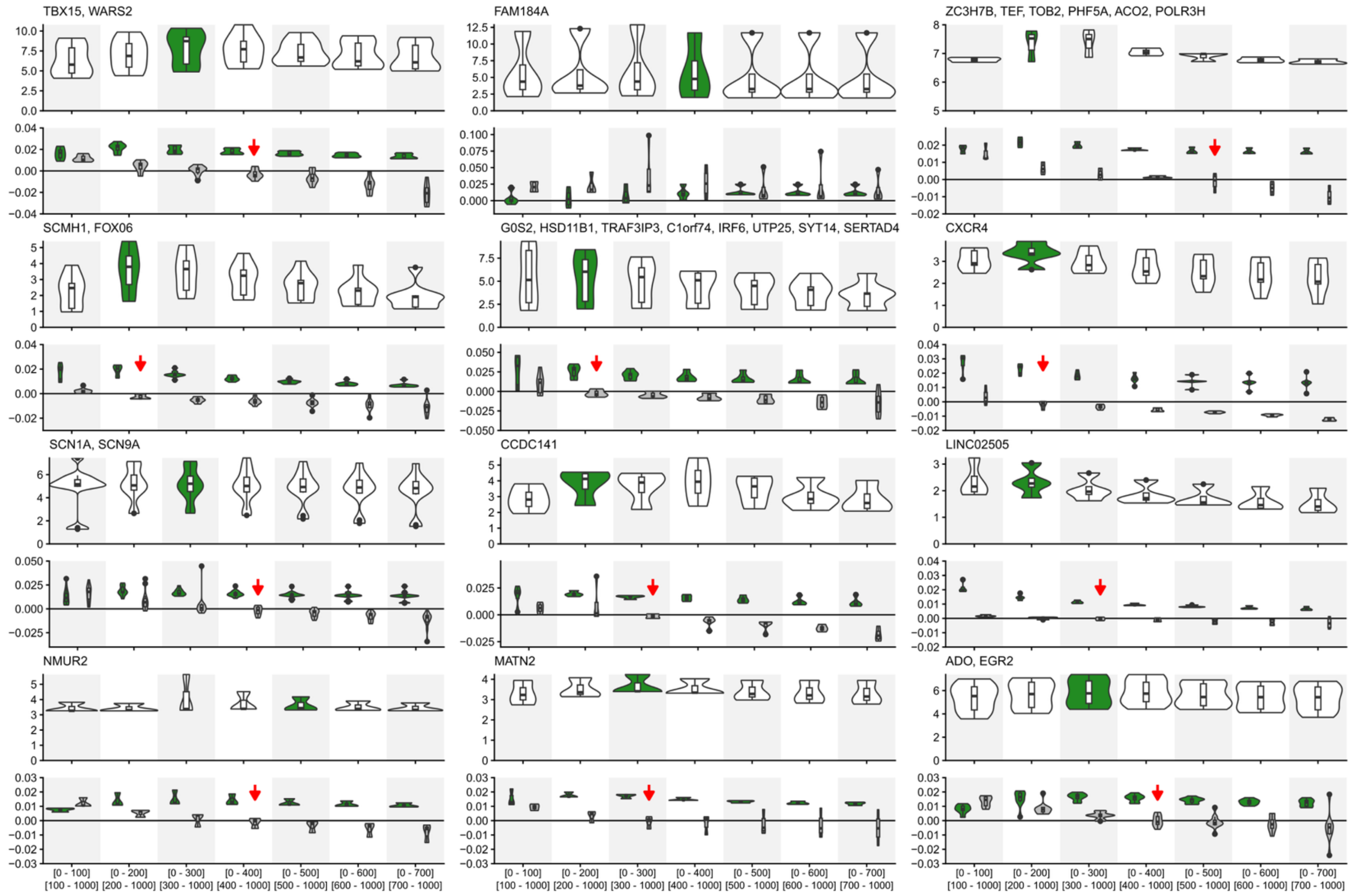


**Fig. S5. Selection History inferred from RELATE ARGs using CLUES2**. Violin plots illustrate the distribution of the inferred logLR for positive selection (top) and selection coefficient (bottom) for each epoch breakpoint (x-axis) across all examined loci. The epoch breakpoint with the highest logLR support for positive selection is colored in green illustrating the best supported epoch breakpoint. The first non-positive epoch 2 is marked with a red arrow illustrating the onset of positive selection coefficients in epoch 2 which we report as the onset of positive selection for each respective locus. ARGs were generated using Relate.

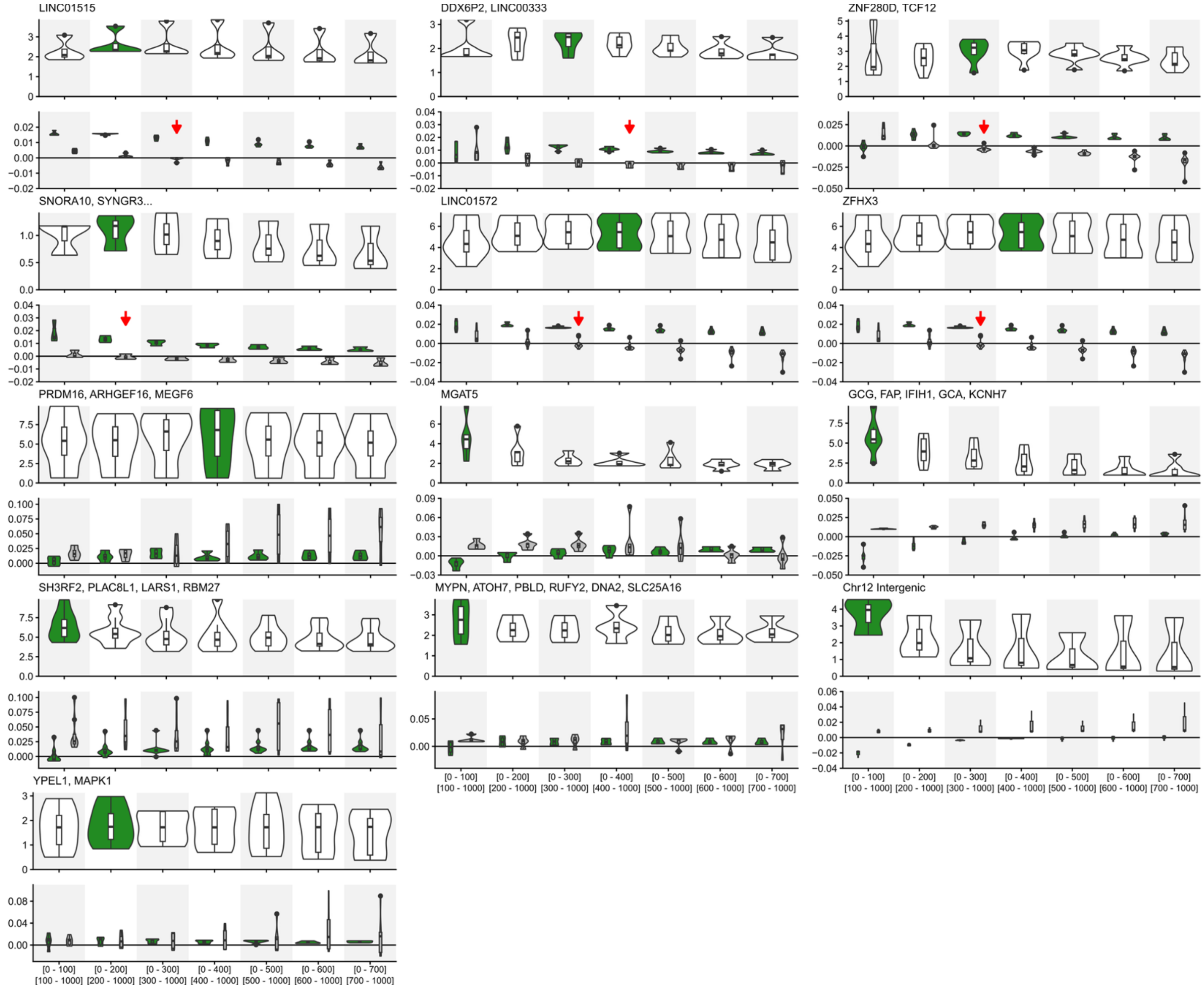


**Fig. S5.** (Continued)

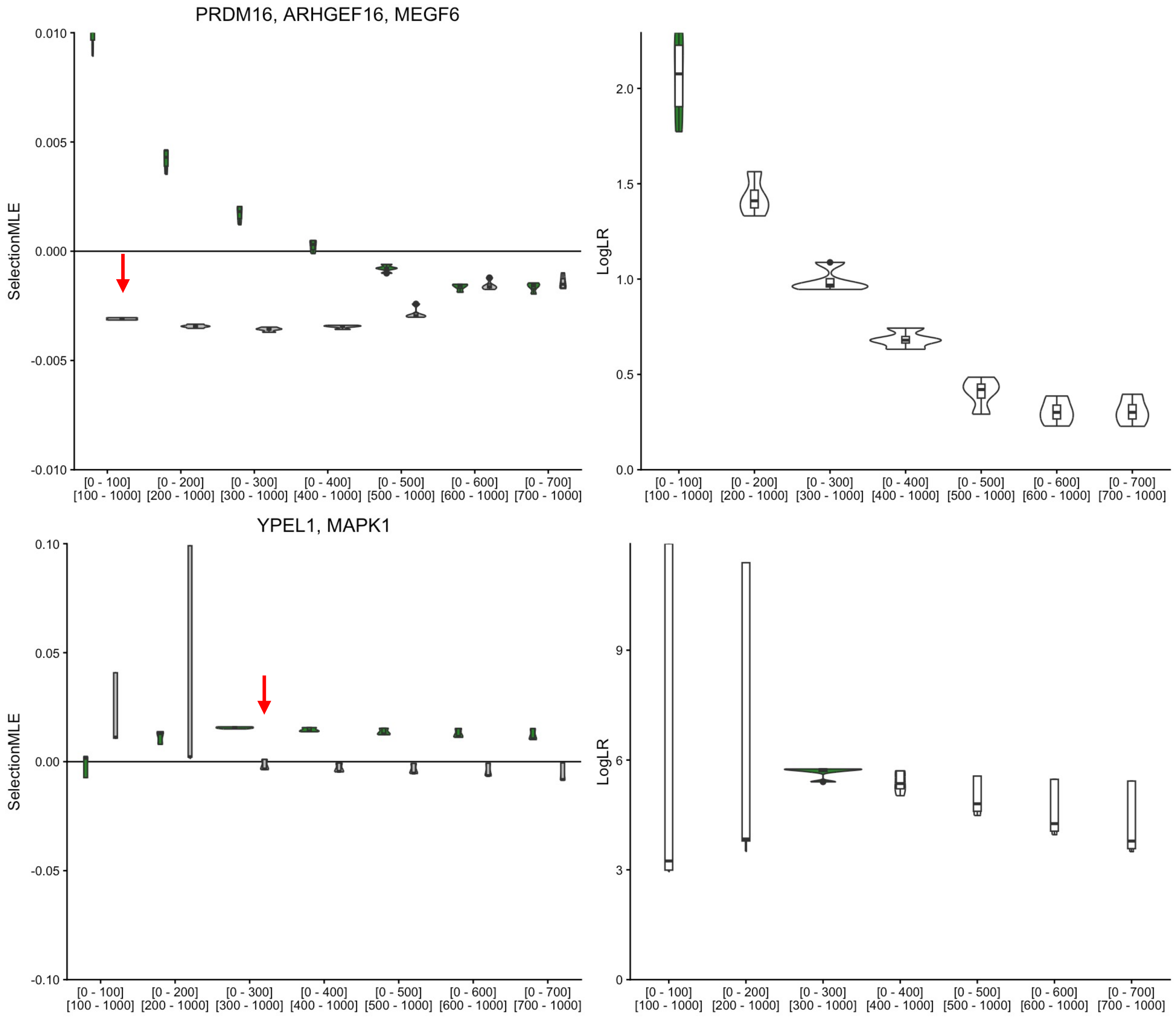


**Fig. S6. Selection History inferred from SINGER ARGs using CLUES2**. Violin plots illustrate the distribution of the inferred logLR for positive selection (top) and selection coefficient (bottom) for each epoch breakpoint (x-axis) across all examined loci. The epoch breakpoint with the highest logLR support for positive selection is colored in green illustrating the best supported epoch breakpoint. The first non-positive epoch 2 is marked with a red arrow illustrating the onset of positive selection coefficients in epoch 2 which we report as the onset of positive selection for each respective locus. ARGs were generated using SINGER.

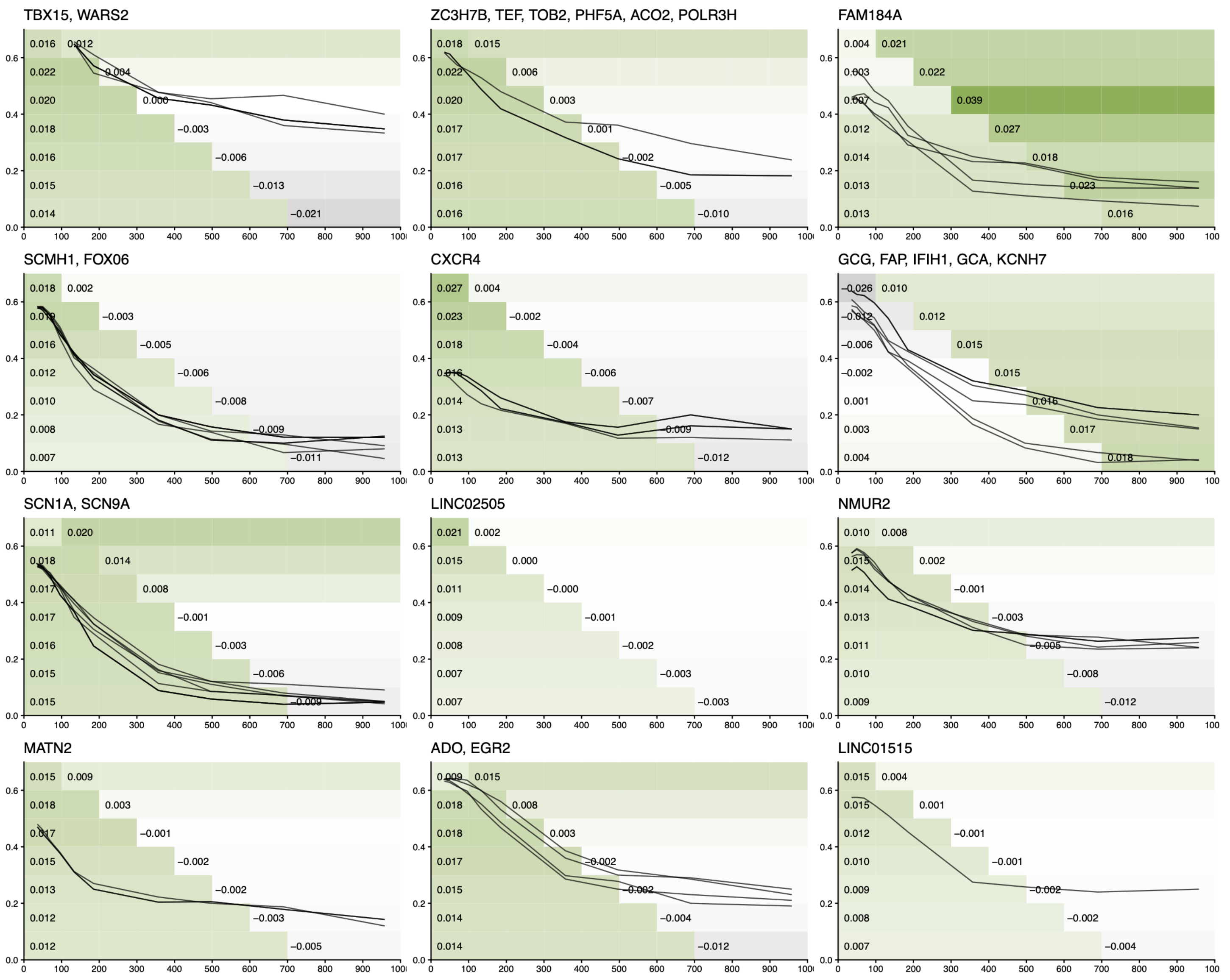


**Fig. S7. Mean selection coefficients across epoch breakpoints and allele frequency trajectories produced using RELATE**. Heatmap: mean CLUES2 selection coefficient (s) inferred under each two-epoch model using RELATE. Each row is a model with a different breakpoint (100 to 700 generations ago, top to bottom). Each column is a 100-generation interval from the present to 1,000 generations ago. Within a row, intervals more recent than the breakpoint show the mean selection coefficient of the recent epoch, and older intervals show that of the older epoch. Coefficients are averaged across the high-frequency U20 alleles used for selection-history inference at this locus. Green indicates positive selection, gray indicates negative selection, and white indicates neutrality. Numeric labels give the mean selection coefficient for the first interval of each epoch. Lines: RELATE-inferred allele frequency trajectories in Peruvians (PEL) for the same U20 alleles.

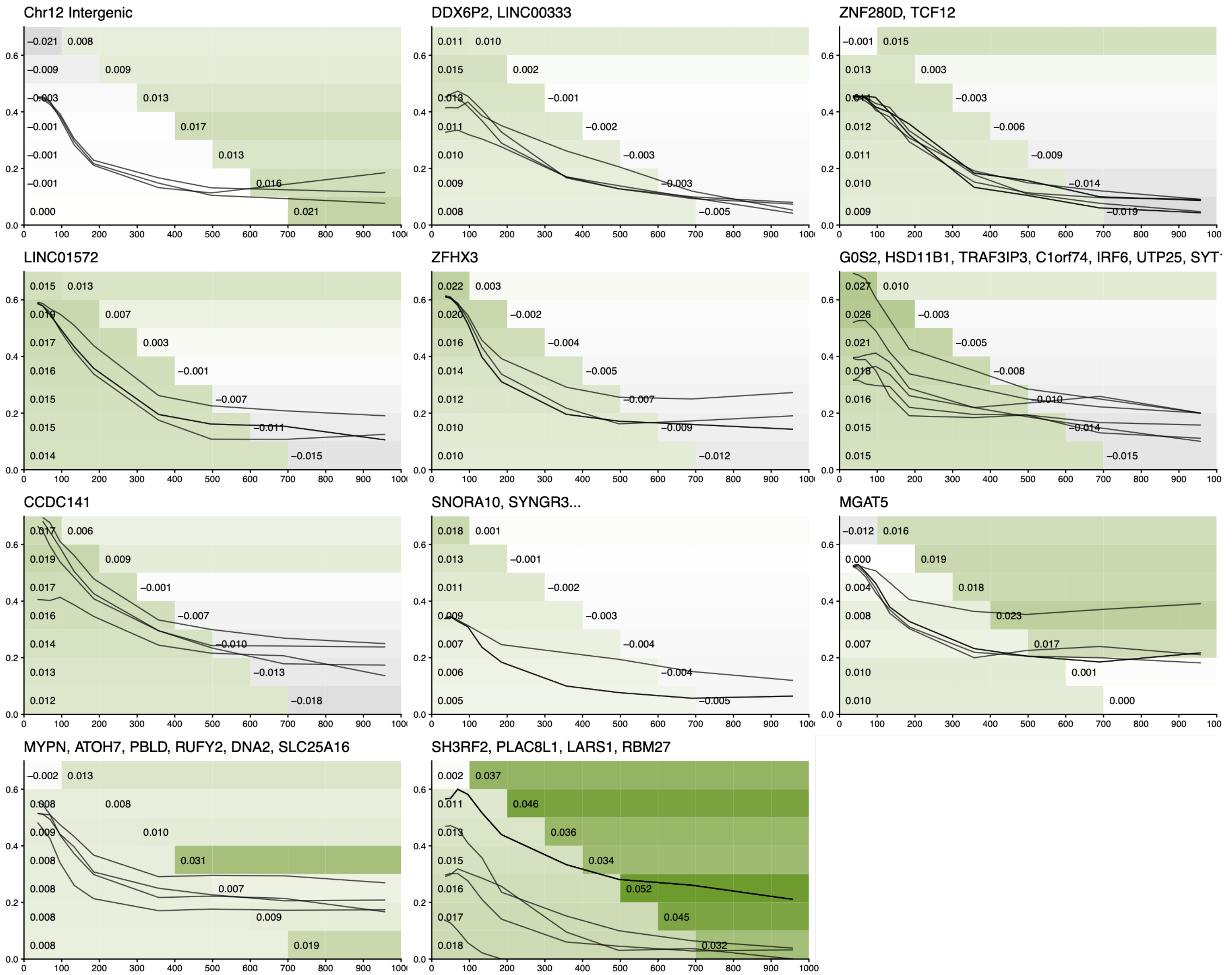


**Fig. S7**. (Continued)

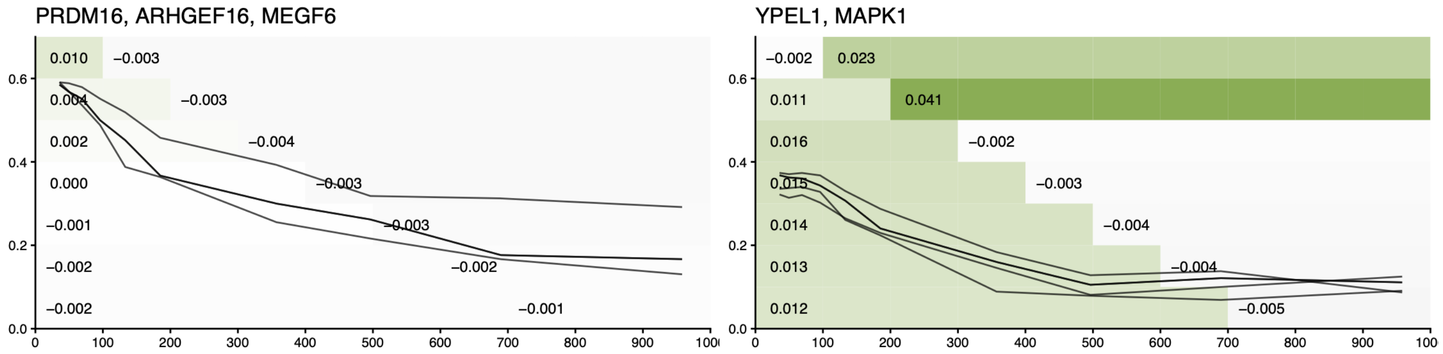


**Fig. S8. Mean selection coefficients across epoch breakpoints and allele frequency trajectories produced using SINGER**. Heatmap: mean CLUES2 selection coefficient (s) inferred under each two-epoch model using SINGER. Each row is a model with a different breakpoint (100 to 700 generations ago, top to bottom). Each column is a 100-generation interval from the present to 1,000 generations ago. Within a row, intervals more recent than the breakpoint show the mean selection coefficient of the recent epoch, and older intervals show that of the older epoch. Coefficients are averaged across the high-frequency U20 alleles used for selection-history inference at this locus. Green indicates positive selection, gray indicates negative selection, and white indicates neutrality. Numeric labels give the mean selection coefficient for the first interval of each epoch. Lines: SINGER-inferred allele frequency trajectories in Peruvians (PEL) for the same U20 alleles.

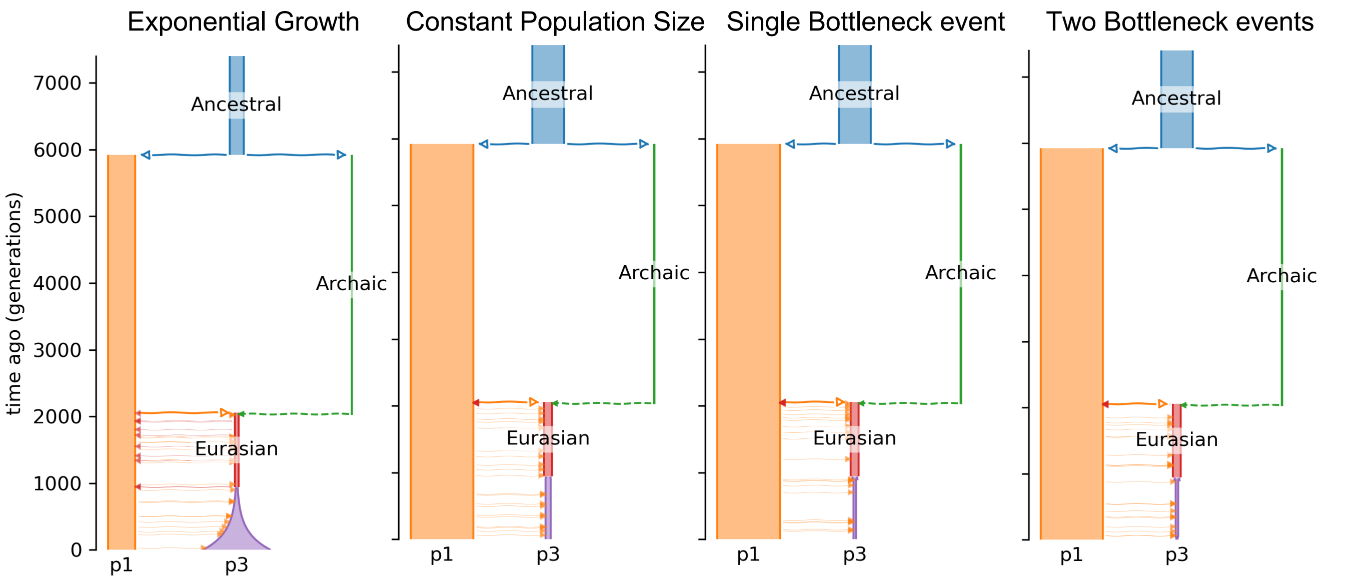


**Fig. S9. Demographic simulation models used to benchmark the selection-inference pipeline**. Simulations included exponential growth, constant population size, a single bottleneck, and two bottleneck events, with modeled relationships among ancestral, archaic, and Eurasian populations. These scenarios were used to assess whether demographic histories relevant to Peruvian populations could confound inference of adaptive introgression and selection onset timing.

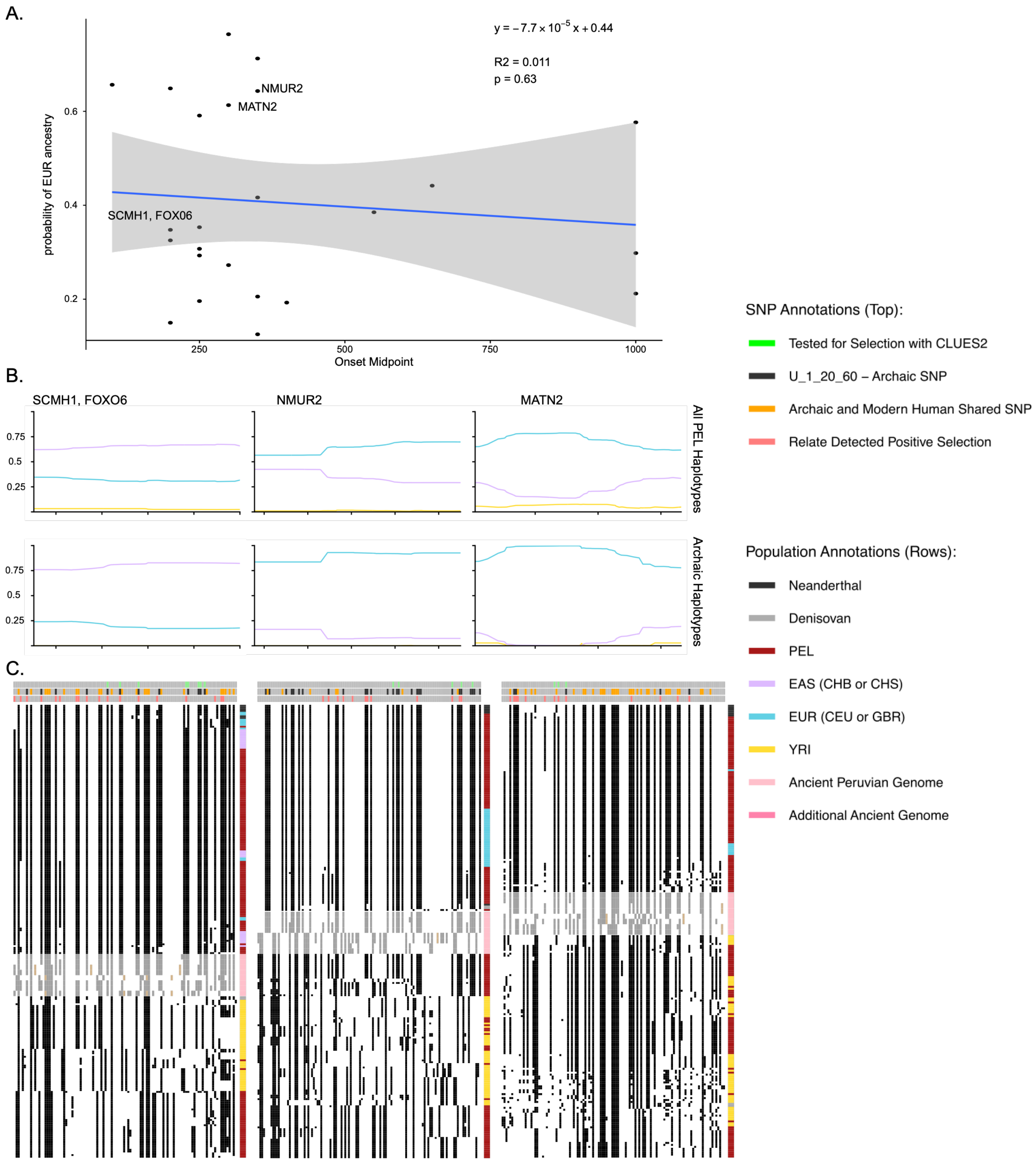


**Fig. S10. Local ancestry is not informative about carrier population, while ancient genomes confirm our inference.** A) Average proportion of European ancestry among all Peruvian haplotypes at each candidate locus (y-axis) against the midpoint of its inferred selection-onset window (x-axis, generations ago) for the 21 dated loci. The blue line is the fitted linear regression, and the gray band is its 95% confidence interval. Labeled loci are shown in B and C. B) Chromopainter local ancestry probabilities across the haplotype region at three candidate loci (*SCMH1*/*FOXO6*, *NMUR2*, *MATN2*).

The top row shows all PEL haplotypes. The bottom row shows PEL haplotypes carrying the archaic-like haplotype, defined as 5% of PEL haplotypes with the smallest distance to the archaic reference. Purple, EAS; blue, EUR; yellow, YRI. The x-axis is genomic position. C) Haplostrip plots for three corresponding candidate regions. Each column is a polymorphic SNP, each row is a single haplotype and the colored column to the right corresponds to the population identity of each haplotype (Black: Neanderthal, Gray: Denisovan, Purple: EAS (CHB or CHS), Blue: EUR (CEU or GBR), Red: PEL, Yellow: YRI, Pink: Ancient Peruvian genome). Gray rows correspond to ancient Peruvian genomes. The dark cells represent the presence of the derived allele and white space represents the ancestral allele.

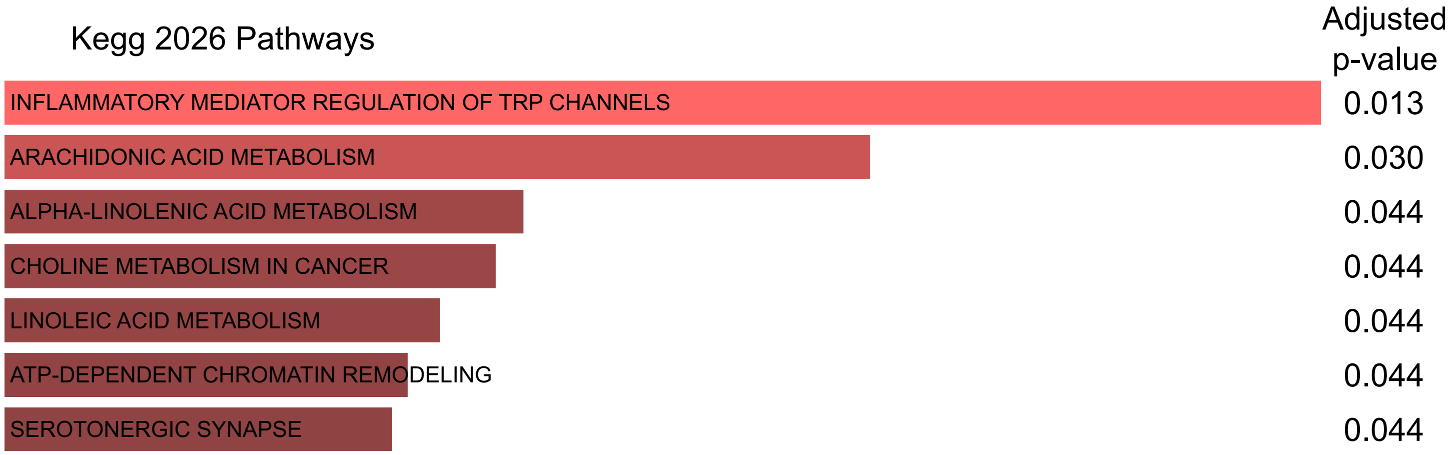


**Fig. S11. Enrichr KEGG enrichment analysis of genes in the top 1% of the genome-wide Q95 distribution**. Adjusted p-values are shows beside each bar with the color of the bar corresponding to the adjusted p-value.

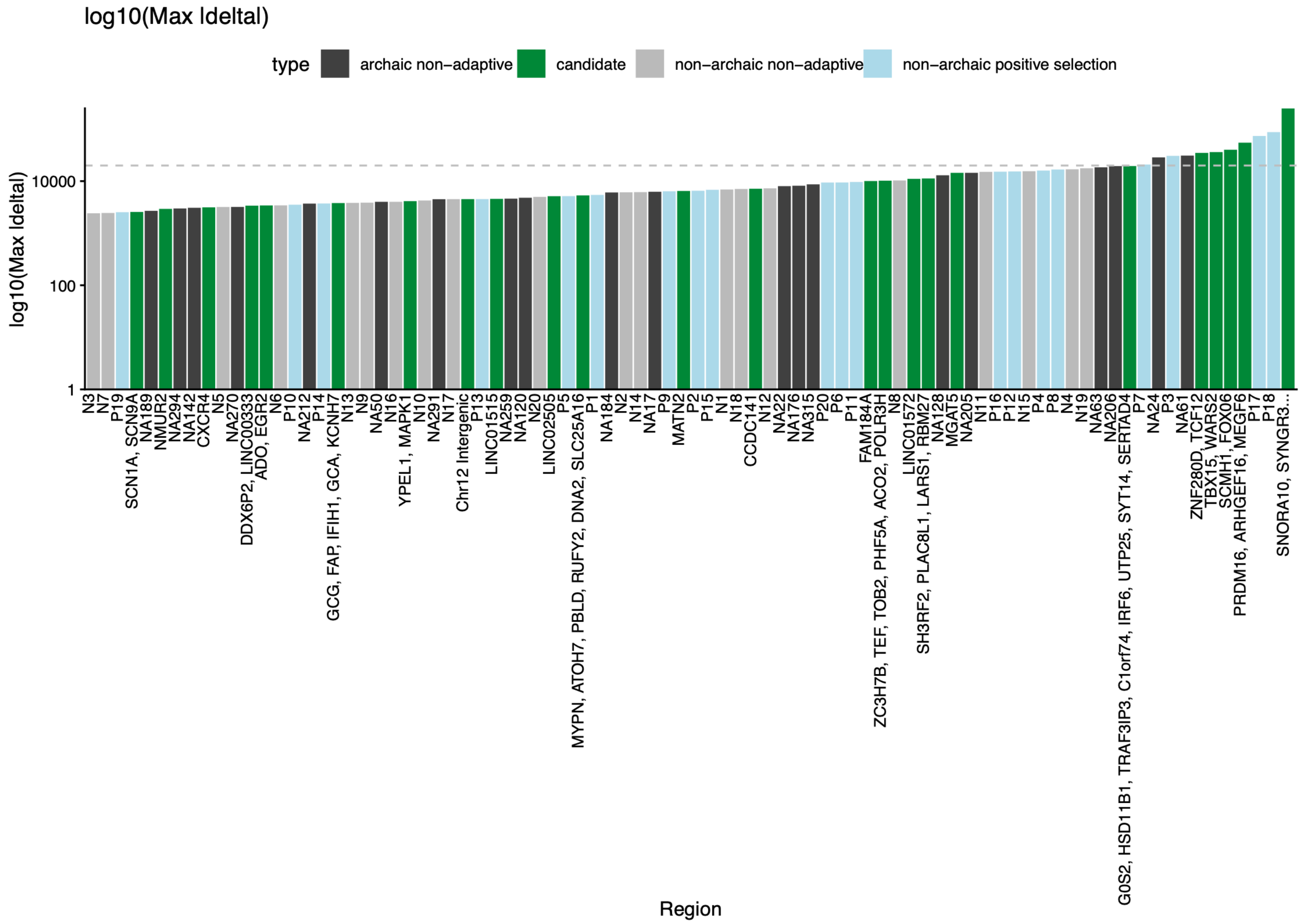


**Fig. S12. Maximum absolute AlphaGenome haplotype delta score, plotted as $\log_{10}(max\,|\Delta|)$, for genomic regions**. Bars are colored by region class: candidate adaptive introgression haplotypes shown in green, archaic non-adaptive regions in dark gray, non-archaic non-adaptive control regions in light gray, and non-archaic regions with evidence of positive selection in light blue. The dashed horizontal line marks the 95th percentile of non-adaptive regions used to highlight regions with elevated predicted regulatory effects.

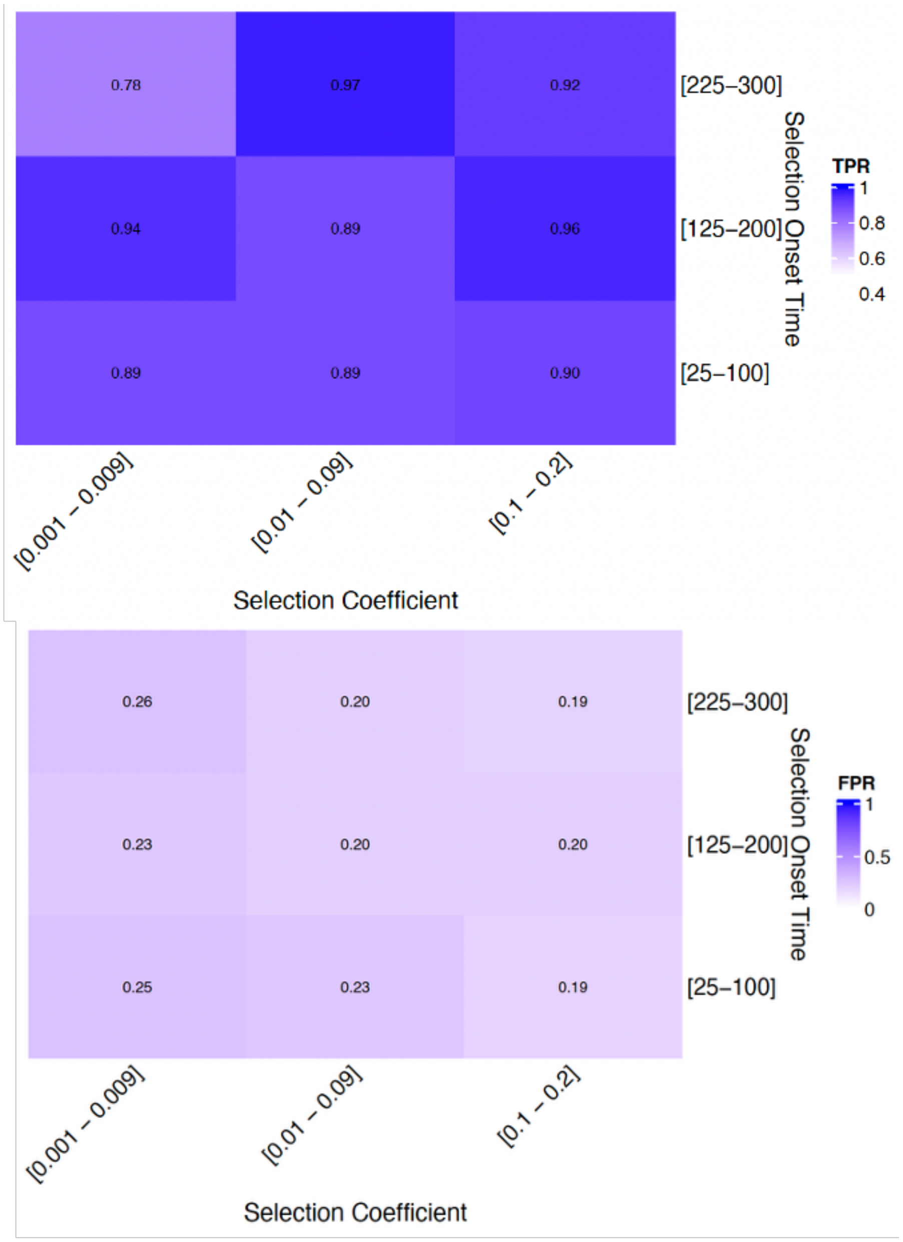


**Fig. S13. Performance of Relate DetectSelection on simulated adaptive introgression.** True positive rates (TPR) and false positive rates (FPR) for detecting selection on introgressed variants in 2-Mb SLiM simulations under the exponential growth demographic scenario in Supplementary Fig. 6. An archaic mutation introduced by a 10% pulse of introgression became beneficial between 25 - 300 generations ago (y-axis) with selection coefficients of s = 0.001–0.02 (x-axis). TPR is the proportion of simulations in which DetectSelection reported a significant U20 variant within 250 kb of the target mutation. FPR is the proportion of non-overlapping 50-kb windows outside this interval that contained at least one significant U20 variant.
